# Translating Innovation to Clinic: End-to-End Bioprocess Development and cGMP Manufacturing of N332-GT5 HIV Vaccine Candidate for First-in-Human Trials HVTN144

**DOI:** 10.64898/2026.06.11.731363

**Authors:** Sammaiah Pallerla, Shaunak Uplekar, Ferenc Boldog, James C. Paulson, Sabyasachi Baboo, John R. Yates, Wen-Hsin Lee, Gabriel Ozorowski, Joel D. Allen, Max Crispin, Chris Cottrell, Andrew B. Ward, Varsha Sitaraman, Tim Broderick, Amber Costakes, Nikolette McCombs, Derek Ryan, Leslie Wolfe, Daniel Craig, Kristen Syvertsen, Albert E. Price, Jon M. Steichen, William R. Schief

## Abstract

The successful translation of rationally designed HIV-1 immunogens into effective vaccines requires manufacturing platforms that maintain structural conformity while meeting clinical-grade quality standards. We developed and scaled a robust, cGMP-compliant process for N332-GT5 gp140, a germline-targeting envelope trimer designed to initiate broadly neutralizing antibody responses, which is now undergoing first-in-human evaluation in HVTN144. Starting with a stable CHO cell line developed using Leap-In® transposon technology, we established a production clone exhibiting high-titer expression (>200 mg/L) and genetic stability through 60 population doublings. The manufacturing process scaled efficiently from Ambr® 250 miniature bioreactors to 200-L single-use systems, delivering consistent product quality across multiple cGMP batches. A streamlined three-step purification strategy—affinity capture, multimodal polishing, and viral clearance- yielded >99% trimeric purity with preserved quaternary structure and native-like antigenicity. Orthogonal LC-MS analyses confirmed site-specific glycan occupancy matching design specifications, while robust viral clearance exceeded 18-log and 11-log reductions for model retroviruses. Clinical material manufactured through this platform has been successfully administered in HVTN144. This work establishes a scalable, reproducible manufacturing paradigm for structurally complex HIV-1 envelope immunogens, advancing the field toward rational vaccine design based on germline-targeting principles.

## 1 Introduction

Developing an effective HIV vaccine continues to be one of the biggest challenges in global public health. Despite over forty years of dedicated research since the HIV/AIDS pandemic began, no vaccine has yet generated the strong, lasting immunity needed to stop HIV infection. With about 1.7 million new cases each year worldwide (UNAIDS, 2025), the urgency to develop a successful preventive HIV vaccine has never been higher.

Traditional vaccine development strategies that have been successful for other pathogens have consistently failed against HIV, mainly due to the virus’s exceptional genetic diversity and its capacity to evade immune responses through rapid mutation and a dense glycan shield on its envelope glycoprotein (Env). The HIV Env trimer is the only target for neutralizing antibodies on the virus surface (Burton et al., 2016), yet it remains one of the biggest immunological challenges in nature. Its high variability, thick glycan shield, and flexible structure have made standard vaccination methods ineffective in producing broadly protective antibodies.

N332-GT5 gp140 represents the culmination of an iterative, structure-guided design process to create an optimal germline-targeting immunogen for BG18-class bnAb precursors. Through multiple rounds of computational design, recombinant protein engineering, and affinity screening against a diverse panel of inferred germline antibodies and next-generation sequencing (NGS)-derived HCDR3 sequences, the N332-GT series (N332-GT1, GT2, and GT5) was progressively optimized, with selection antibodies used at each stage to isolate the highest-affinity clones and incorporate the best mutations into the next-generation Env immunogen (Steichen et al., 2019). N332-GT5 emerged as a highly refined immunogen that binds 20 of 27 potential BG18 precursor antibodies with Kd ≤ 1 μM, meeting the critical threshold required for robust germinal center activation and B cell priming.

Preclinical studies in rhesus macaques subsequently demonstrated the remarkable efficacy of N332-GT5 when formulated with the saponin/MPLA nanoparticle (SMNP) adjuvant (Steichen et al., 2024). Immunization with N332-GT5/SMNP successfully primed BG18-class precursor B cells in all eight animals tested — the first demonstration of priming of HCDR3-dominant bnAb precursors in outbred animals. Cryo-electron microscopy structural analysis confirmed that vaccine-elicited antibodies bound to HIV Env trimers with the characteristic BG18-class binding footprint, angle of approach, and HCDR3-dominant interaction. Immunization also induced robust affinity maturation, with binding affinities improving over 6,000-fold within 10 weeks — from a median Kd of 98 nM to less than 4 pM — and antibodies gaining cross-reactivity to more native-like HIV Env trimers. Furthermore, 65% of BG18 type I antibodies isolated at weeks 7 and 10 post-prime could bind to at least one trimer containing all glycans in the N332 epitope, a necessary requirement for an N332-dependent bnAb, and demonstrated the ability to neutralize pseudoviruses expressing BG505-based envelope proteins.

Based on these highly promising preclinical results, N332-GT5 gp140 advanced to clinical evaluation in the HVTN 144 phase 1 clinical trial. HVTN 144 is the first-in-human evaluation of this germline-targeting immunogen and tests multiple vaccination strategies, including different doses, routes of administration (intramuscular versus subcutaneous), and dosing schedules (bolus versus fractionated escalating-dose prime). The study aims to evaluate the safety and tolerability of N332-GT5 gp140 adjuvanted with SMNP (Silva et al., 2021; Pallerla et al., 2025) and, critically, to assess whether this immunogen can successfully prime BG18-class B cell responses in humans, as it did in non-human primates. HVTN 144 is also the first human trial to test a generalizable strategy for priming HCDR3-dominant bnAbs, with implications beyond HIV for other pathogens.

Native-like soluble Env trimers like BG505 SOSIP.664 have enabled the induction of neutralizing antibody responses in preclinical models (Sanders et al., 2015). Subsequent engineering strategies improved trimer stability and facilitated germline-targeting immunogen design (Steichen et al., 2016). More recently, clinical evaluations have demonstrated the safety and immunogenicity of native-like trimer vaccines in humans (Parks et al., 2025).

The successful translation of N332-GT5 gp140 from concept to clinical-grade material required overcoming significant process development and manufacturing hurdles. Producing native-like HIV Env trimers at the scale, purity, and consistency needed for clinical trials involved establishing reliable, reproducible bioprocessing platforms that maintain the complex structural and antigenic properties essential for immunogen function.

This manuscript details the complete development and manufacturing process of N332-GT5 gp140 for clinical use in HVTN144. It emphasizes the strategic decisions, technical advancements, and quality-control procedures implemented to produce clinical-grade material that complies with strict regulatory standards for Phase I human trials while preserving the key structural and immunological characteristics of this next-generation HIV vaccine candidate.

## 2 Materials and Methods

### 2.1 Analytical Methods-Cell Line Development

#### 2.1.1 SDS PAGE

For reduced SDS-PAGE gels, clarified supernatant was prepared with NuPAGE™ LDS Sample Buffer (4X) (Invitrogen, Catalog#: NP0008) and reduced using NuPAGE™ Sample Reducing Agent (10X) (Catalog#: NP0009) following the vendor protocol. The reduced samples were run on NuPAGE Novex 4-12 Bis-Tris Protein Gels (Invirogen, Catalog#: NP0329BOX, WG1403BX10) and stained with InstantBlue™ Protein Stain (Invitrogen, Catalog#: ISB1L-1L) according to the manufacturer’s instructions. For non-reduced SDS-PAGE gels, clarified supernatant was prepared with NuPAGE™ LDS Sample Buffer (4X) (Catalog#: NP0008) as per the manufacturer’s protocol. Additionally, 10 mM N-Ethylmaleimide (NEM) (Catalog#: E3876-5G) was added to prevent reformation of disulfide bonds. The samples were then run on NuPAGE Novex 4-12 Bis-Tris Protein Gels (Invitrogen, Catalog #: NP0329BOX, WG1403BX10) and stained with InstantBlue™ Protein Stain (Catalog #: ISB1L-1L) according to the vendor’s instructions.

#### 2.1.2 Octet BLI binding assay

Trimer titer was assessed on a ForteBio Octet RED384. Detection antibodies (positive: PGT145 antibody (directed against a quaternary structure located at the trimer apex), BG18_GL0 (directed against the glycan-V3 portion of Env) antibody, negative: DEN3 antibody, (Steichen et al., 2019) were diluted to 10 µg/mL in 1X Assay Buffer [PBS pH 7.4, 1:10 vol of 10X Kinetics Buffer (ForteBio Cat. 18-1105)]. IgGs were captured on Anti-Human Fc Capture (AHC, ForteBio Cat. 18-5060) biosensor tips. Loaded tips were equilibrated in 1X Assay Buffer with 5% sterile custom EX-CELL Advanced CHO Fed-batch (AFB) media (SAFC, St. Louis, MO), then transferred to clarified supernatant samples diluted 1:20 in 1X Assay Buffer. The initial binding rate V0 was determined by subtracting the negative DEN3 trace from the positive PGT145 or BG18_GL0 antibody signal. V0 values were converted to µg/ml using a standard curve of purified N332-GT5 gp140, diluted in 1X Assay Buffer with 5% sterile AFB media. Between measurements, the antibody and remaining N332-GT5gp140 trimer were stripped from the tips by regeneration in 10 mM Glycine pH 1.7.

#### 2.1.3 Transcript sequence analysis

Total RNA was isolated from the final C235 clone cell pellets (1×10^6^ cells/sample) using the RNeasy Mini Kit (Qiagen). Complementary DNA fragments corresponding to the region encoding N332-GT5 gp140, or human furin were amplified using the OneStep RT-PCR Amplification Kit (Qiagen Cat. 210212). The amplified products were sequenced using gene-specific primers and Sanger chemistry with 100% double-stranded coverage. The presence and the perfect match to the expected sequences have been confirmed for both recombinant messages.

### 2.2 Analytical Methods-Process Development and Manufacturing

The non-compendial analytical methods used to characterize and control the N332-GT gp140 drug substance are outlined below. Methods for testing appearance, pH, endotoxin, and bioburden, which are covered by compendial procedures, were performed according to established pharmacopeial standards and are therefore not included.

#### 2.2.1 Determination of Protein Concentration by Absorbance at 280 nm

Protein concentration was measured using a SoloVPE variable pathlength spectrophotometer (C Technologies, Inc., Cedar Knolls, NJ) based on UV absorbance at 280 nm. The extinction coefficient (ε) of 1.686 mL·mg⁻¹·cm⁻¹, calculated from the trimeric amino acid sequence, was used according to the Beer–Lambert law.

Concentration (mg/mL) was determined by:

C = A_280_ − A_320_ / 1.686
where A_280_ and A_320_ are the absorbance values at 280 nm and 320 nm, respectively.

#### 2.2.2 Size Exclusion High Performance Liquid Chromatography (SE-HPLC)

Protein purity and aggregation state were assessed through size exclusion chromatography using a TSKgel UltraSW Aggregate column (TOSOH, Cat#22856). Samples were injected to reach a target column load of 15µg of N332-GT gp140 drug substance and subsequently examined under isocratic conditions containing phosphate buffer as the mobile phase. Separation was detected at 280 nm to measure monomeric, high-molecular-weight (HMW), and low-molecular-weight (LMW) species.

#### 2.2.3 Quantification of Residual CHO DNA by qPCR

Residual host cell DNA was quantified by real-time quantitative PCR (qPCR) using a Bio-Rad CFX384 Real-Time PCR Detection System (Bio-Rad Laboratories). The assay used CHO-specific primers and a TaqMan® probe targeting a conserved genomic sequence. The fluorescence signal produced during amplification was analyzed with CFX Manager software. DNA concentration was determined by comparing it with a standard curve generated from known concentrations of CHO DNA.

#### 2.2.4 Quantification of Host Cell Proteins (HCP)

Residual CHO host cell proteins were measured using a third-generation CHO HCP ELISA kit (Cygnus Technologies, Southport, NC). The two-site immunoenzymatic assay employed affinity-purified anti-CHO polyclonal antibodies—one coated on the microtiter plate and the other conjugated to horseradish peroxidase (HRP). After incubation and washing steps, 3,3’,5,5’-tetramethylbenzidine (TMB) substrate was added, and absorbance was read at 450 nm. HCP concentrations were determined from a standard curve fitted to a four-parameter logistic (4PL) equation.

#### 2.2.5 Biolayer Interferometry (BLI)

Biolayer interferometry (BLI) was utilized to examine biomolecular interactions and binding activity of N332-GT gp140. The assay was conducted with a FortéBio Octet system (Sartorius, Fremont, CA). This label-free optical method monitors real-time shifts in the interference pattern caused by analyte molecules binding to immobilized ligands on biosensor tips. Association and dissociation kinetics, binding specificity, and relative concentrations were assessed by fitting data to appropriate kinetic models.

#### 2.2.6 Purity by Reverse Phase Chromatography

Protein purity and identity were assessed through reverse phase chromatography using an Acquity UPLC Protein BEH C4 300Å column (Cat# 186004497, Waters Corporation, Milford, MA). Samples were injected to reach a target column load of 6.0 µg of N332-GT gp140 drug substance and was examined over a gradient generated by mobile phases composed of TFA, acetonitrile and water. Separation was detected at 214 nm to measure pre-Main, Main and post-Main species.

#### 2.2.7 Quantification of Residual 2G12

Residual IgG levels were quantified using a Protein A–based ELISA. Samples were captured on Protein A–coated plates, detected using an alkaline phosphatase–conjugated secondary antibody, and developed with p-nitrophenyl phosphate substrate prior to absorbance measurement at 405 nm. IgG (2G12) concentrations were calculated by interpolation from a standard curve generated using reference standards.

#### 2.2.8 Determination of Residual Triton X-100

Residual Triton X-100 was quantified using a ZORBAX StableBond 300 C8 column (Cat.# 865750-906, Agilent, Santa Clara, USA). Samples were injected with a 10µL volume and was examined over a linear gradient generated by mobile phases composed of acetonitrile and water with detection at 276 nm.

#### 2.2.9 Determination of Residual Protein A

Residual Protein A levels were quantified using the Mix-N-Go Protein A ELISA Kit with Amsphere™ A3 (JSR Life Sciences, CA, USA) Protein A standards (F610, Cygnus Technologies). Samples were captured on Anti-Protein A–coated plates, detected using a horse radish peroxidase–conjugated secondary antibody, and developed with 3,3’,5,5’-tetramethylbenzidine (TMB) substrate prior to absorbance measurement at 450 nm and 650nm. Residual Protein A concentrations were calculated by interpolation from a standard curve generated using Protein A standards.

### 2.3 Cell Line Development

The Cell line development workflow for stable pools and clone generation is shown in Figure 1.

**Figure 1:**
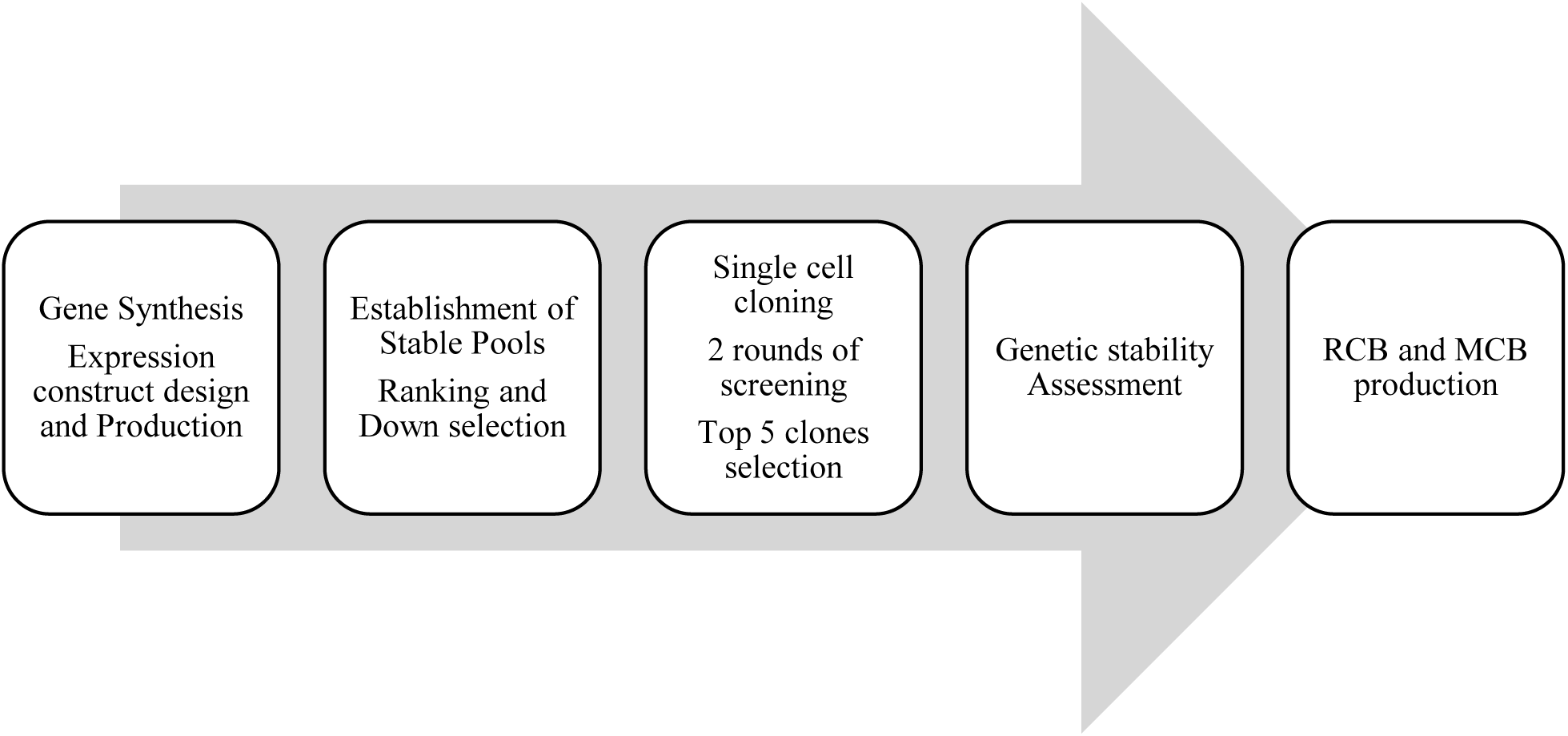
Workflow for Stable Pools and clone generation

#### 2.3.1 Parental Cell Line

HD BIOP3 is a GS-null cell line derived from ECACC CHOK1 established by Horizon Discovery.

#### 2.3.2 Leap-In® Transposase System

The Leap-In ® transposase system, composed of five different transposons and their corresponding cognate transposases, has been developed by ATUM. The enzymes belong to the DD[D/E] integrase family and catalyze a cut-and-paste mechanism to insert their cognate transposons into the target genome. The enzymatic mechanism results in single-copy integrants at multiple genomic loci. The transposase-mediated integration significantly reduces the formation of concatemers and the deletion and rearrangement of transgenes. Leap-In mediated transposition increases the likelihood of integration into transcriptionally active genomic regions.

#### 2.3.3 Coding Sequences, Expression Vector Construction

Two expression constructs were designed in the Leap In1 transposon-based backbone. One construct expressed the N332-GT5 gp140 codon-optimized coding sequence, and the other expressed a codon-optimized human furin ORF. The codons were optimized using ATUM’s proprietary algorithm. Both constructs expressed a glutamine synthetase cassette. All DNA sequences were chemically synthesized from phosphoramidites, cloned into intermediate vectors, and transformed into *E. coli* hosts using ATUM’s proprietary technology. Transfection quality midi-preps were made from the constructs, and their sequences were confirmed by Sanger sequencing.

#### 2.3.4 Stable Pool Development

HD BIOP3 cells were transfected by electroporation with 25 µg of 3:1 ratio of N332 to furin plasmid DNA and 3 µg Leap In 1 mRNA. A stable pool was established in glutamine-free selection medium. At viability >95%, the pool was cryopreserved.

#### 2.3.5 Fed Batch Production

Productivity and product quality were assessed using material produced in fed batch production runs. The runs were performed in Advanced Fed Batch media, in either 250 mL shake flasks (50 mL working volume), Tube spin reactors (10 mL working volume) or 24 deep well plates (3 mL working volume) using ATUM’s proprietary culture conditions using Cellboost 7, Advanced CHO Feed, and Cellvento 4 Feed. The 250 mL shake flask and Tubespin scale cultures were sampled on days 7, 10, 12, and harvested on day 14, while the 24 deep well cultures were sampled on day 5 and harvested on day 7. The harvests were clarified by centrifugation and filtration through 0.2 µm membranes.

#### 2.3.6 Single Cell Cloning

Clonally derived N332-GT5 gp140 producing cell lines were established by performing one round of single cell cloning using the VIPS single clone deposition and imaging instrument. Clonal cell line deposition and verification of monoclonality were performed using the VIPS and Cell Metric systems (Solentim/Advanced Instruments).

#### 2.3.7 Clone Ranking

##### 2.3.7.1 First Round of Clone Ranking

The first clone ranking was executed in two stages. In both cases, seven-day fed-batch runs were performed at a 24-deep-well scale. Following a BG18_GL0 detectable productivity assessment on 213 clones by BLI/ Octet (Pall) instrument, a set of 56 clones was selected to establish the BG18_GL0/PGT145 detectable productivity ratios. Based on these ratios and the clonal growth characteristics, twenty-four clones were selected for the second round of clone ranking.

##### 2.3.7.2 Second Round of Clone Ranking

Fourteen-day Tubespin scale-fed batch runs were initiated for the selected 24 clones. The ranking parameters were the clonal BG18/PGT145 detectable productivity ratio, the efficiency of furin cleavage assessed by reduced SDS PAGE, and clonal growth characteristics.

#### 2.3.8 Confirmation of Clonal Genetic Stability

Sixty population doubling (PD) long serial passage cultures were maintained in 24 deep well plates, in the absence and in the presence of 5 mM glutamine. PD0, PD60+Gln, and PD60-Gln cells were used to initiate 12-day fed batch runs in TubeSpins (10 mL culture). Productivity, transgene copy numbers, and N332 and furin mRNA identities were used as stability indicators.

### 2.4 Cell Culture Process Development

The upstream cell culture process development (Figure 2) began with the selected research cell bank (RCB) of clone C235. The development strategy included key steps such as media adaptation, process optimization in a high-throughput Ambr^®^250 bioreactors, and scale-up to a pilot scale bioreactor. Verification and demonstration of process performance was performed across multiple pilot scale bioreactors, using both the RCB and MCB for the selected clone. Media adaptation involved transitioning the clone from EX-CELL^®^ Advanced™ fed-batch media (SAFC, St. Louis, MO) to commercially available Dynamis™ (Thermofisher Scientific, Waltham, MA) media over five passages by direct adaptation. Following successful adaptation, process optimization was conducted using the Ambr®250 (Sartorius, Gottingen, Germany) high-throughput system to identify the optimal process parameters, including media composition and culture conditions, that maximize cell growth and productivity. Once the parameters were finalized and verified in the Ambr®250 system, the process was scaled up to Xcellerex^™^ XDR-200 bioreactor (Cytiva, Marlborough, MA). Initial runs utilized the RCB, with later runs incorporating both the RCB and the MCB to ensure robustness across cell lines. This scale-up established the final upstream cell culture process for producing GMP-grade material for clinical trials, using Xcellerex™ XDR-200 (Cytiva, Marlborough, MA) in GMP suites.

**Figure 2:**
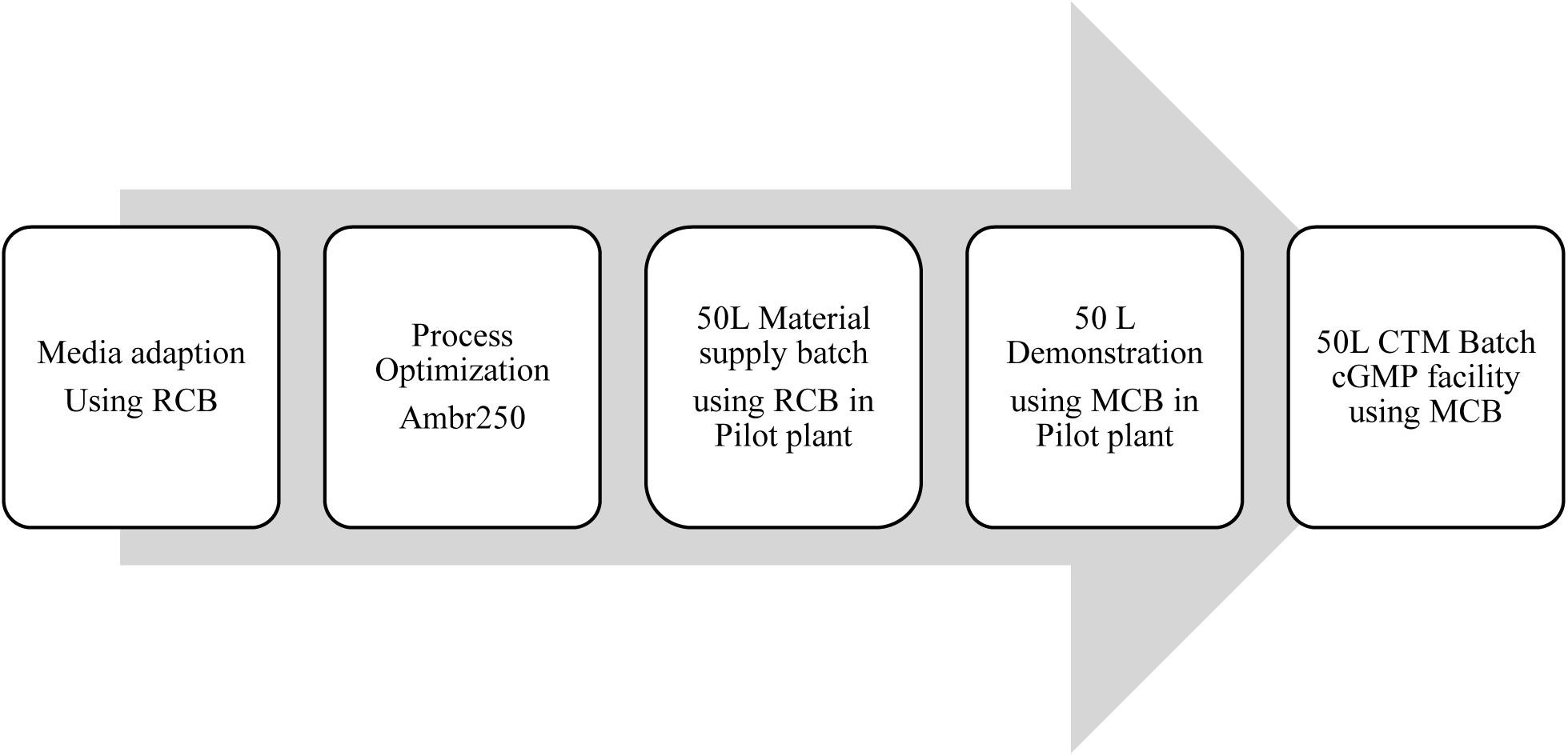
Workflow for Cell Culture Process Evaluation

#### 2.4.1 Cell Line and Medium Adaptation

A research cell bank (RCB) of CHO clone C235 was obtained from the cell line development partner (ATUM) supplied in custom EX-CELL Advanced CHO Fed-batch (AFB) medium (SAFC, St. Louis, MO) without dextran sulfate. Due to limited availability and procurement constraints associated with this custom formulation, clone C235 was directly adapted to the commercially available dextran-sulfate-free Dynamis medium (Thermofisher Scientific, Waltham, MA). Dextarn-sulfate free medium was necessary for the process due to the interference of dextran sulfate with product titer measurement using biolayer interferometry (BLI).

Direct adaptation was performed over five consecutive passages, starting from thawed RCB vials. The initial recovery passage (passage 1) was carried out in 125 mL shake flasks using EX-CELL AFB medium. Beginning with passage 2, cells were cultured exclusively in Dynamis medium. The cultures were expanded stepwise in 250 mL and 1 L shake flasks before being scaled up to 5 L flasks by passage 5. Each passage was maintained for approximately 3 days at target inoculum densities of 0.3–0.5 × 10^6 cells/mL under standard conditions (37 °C, 5% CO₂, 120 rpm). Passage 5 material, demonstrating stable growth and viability in Dynamis medium, was subsequently used to generate eight 1 mL research cell bank vials and to inoculate a 12 x Ambr^®^250 process optimization study.

#### 2.4.2 Process Optimization using Ambr®250 bioreactor

The cell culture from medium adaptation was used to inoculate 12 x Ambr^®^250 bioreactors for the process optimization study with clone C235. Ambr^®^250 vessels have been shown to provide scalable data for fed batch production processes (Rameez et al., 2014). The study parameters for the process optimization are outlined in Table 1. The Ambr^®^250 working volumes started from 190 mL on Day 0 to 250 mL on Day 14. The Cell Boost 7a (Cytiva, Marlborough, MA) and Cell Boost 7b (Cytiva, Marlborough, MA) feeds were added to the culture from Day 3 to Day 13 to maintain the nutrient levels as shown in Table 1.

**Table 1:** Study parameters for ambr250 Process Optimization.

| # | Study Factor | Test Parameter Details | Trend Name |
| --- | --- | --- | --- |
| 1 | Control #1 | 27% CB7a slow, 2.7% CB7b bolus, 34°C Day 4 | Control #1 |
| 2 | Control #2 | 27% CB7a slow, 2.7% CB7b bolus, 34°C Day 4 | Control #2 |
| 3 | Control + 7a bolus | 27% CB7a bolus, 2.7% CB7b bolus, 34°C Day 4 | Control + 7a bolus |
| 4 | 35% 7a slow, 3.5% 7b bolus | 35% CB7a slow, 3.5% CB7b bolus, 34°C Day 4 | 35% 7a slow |
| 5 | 35% 7a bolus, 3.5% 7b bolus | 35% CB7a bolus, 3.5% CB7b bolus, 34°C Day 4 | 35% 7a bolus |
| 6 | Inoc @ 1.0e6,<br>40% 7a slow, 4% 7b bolus | Inoc bioreactor to 1.0 e6 cells/mL,<br>40% CB7a slow + 4% CB7b bolus Days 2-13, 34°C Day 4 | High inoc,<br>40% 7a slow |
| 7 | Inoc @ 1.0e6,<br>40% 7a bolus, 4% 7b bolus | Inoc bioreactor to 1.0 e6 cells/mL,<br>40% CB7a bolus + 4% CB7b bolus Days 2-13, 34°C Day 4 | High inoc,<br>40% 7a bolus |
| 8 | Inoc @ 1.0e6,<br>40% 7a slow, 4% 7b bolus,<br>34°C Day 5 | Inoc bioreactor to 1.0 e6 cells/mL,<br>40% CB7a slow + 4% CB7b bolus Days 2-13,<br>34°C Day 5 | High inoc,<br>40% 7a slow,<br>34°C Day 5 |
| 9 | Control + 34°C Day 5 | Control + 34°C Day 5 | 34°C Day 5 |
| 10 | pH 6.90 ± 0.10 | pH setpoint = 6.90 ± 0.10 | pH 6.90 |
| 11 | pH 6.90 + pH 7.20 shift on Day 6 | pH = 6.90 ± 0.10 (Days 0-5), pH = 7.20 ± 0.20 (Days 6-14) | pH 6.90,<br>pH 7.20, Day 6 |
| 12 | pH 7.00 + pH 7.20 shift on Day 6 | pH = 7.00 ± 0.20 (Days 0-5), pH = 7.20 ± 0.20 (Days 6-14) | pH 7.00,<br>pH 7.20, Day 6 |

#### 2.4.3 Pilot and GMP scale runs

A 50L material supply run was conducted in a Cytiva XDR50 Single-Use Bioreactor (Cytiva, Marlborough, MA) (SUB) using the ‘Control + 7a bolus’ condition from the ambr250 process optimization and a C235 RCB to supply clarified harvest material for downstream development studies and to generate KBI Reference Material for N332-GT5 gp140. Subsequently, a 50 L demonstration run was conducted in a Cytiva XDR200 Single-Use Bioreactor (SUB) using the same production process and a C235 Master Cell Bank (MCB) to supply clarified harvest material for downstream process demonstration studies. The seed train for both the runs involved 5 passages in a shake flask and a 6^th^ passage in a wave bioreactor (Cytiva, Marlborough, MA) as an N-1 stage. The wave culture was used to inoculate the production bioreactor. Process conditions were aligned as closely as possible between the two studies to enable direct comparability. Both bioreactors were operated at 37 °C, with a temperature shift to 34 °C on Day 4, a pH setpoint of 7.00 ± 0.20, and a dissolved oxygen (DO) level of 40%. Cell Boost 7a and Cell Boost 7b were added as boluses according to predefined feed schedules. Antifoam (10% ADCF, Thermofisher Scientific, Waltham, MA) and 1 M sodium carbonate (SAFC, St. Louis, MO) were used to control foam and pH. Harvest was performed using D0HC (MilliporeSigma, MA, USA) and X0HC (MilliporeSigma, MA, USA) depth filters on Day 14 or earlier if culture viability dropped below 70%. Table 2 summarizes the bioreactor operating parameters for the two pilot scale production runs. Only the parameters that differed between scale are shown.

**Table 2:** Comparison of process parameters for the 50-L RCB and MCB production runs.

| Process Parameter | 50-L Material Supply (RCB) – XDR-50 | 50-L Demonstration (MCB) – XDR-200 (only differences shown) | 50-L GMP (MCB) – XDR-200 (only differences shown) |
| --- | --- | --- | --- |
| Vessel | GE/Cytiva XDR-50 SUB | GE/Cytiva XDR-200 SUB | GE/Cytiva XDR-200 SUB |
| Day 0 Weight | 34.0 ± 0.5 kg | 40.0 ± 0.5 kg | 50.0 ± 0.5 kg |
| Medium Hold Weight | 29.4 kg Dynamis + 0.6 kg L-Gln | 39.2 kg Dynamis + 0.8 kg L-Gln | 50.5 kg Dynamis + 1.0 kg L-Gln |
| Temperature Setpoint | 37 °C → 34 °C (Day 4) | Same | Same |
| pH Setpoint | 7.00 ± 0.20 | Same | Same |
| DO Setpoint | 40 % | Same | Same |
| Feed Type | CB7a and CB7b (bolus) | Same | Same |
| Glucose and Feed Targets | Same | Same | Same |
| Antifoam / Base | 10% ADCF / 1 M sodium carbonate | Same | Same |
| Harvest Criteria | Day 14 or <70% viability | Same | Same |

### 2.5 Downstream Purification Process Development

The downstream purification process for N332-GT5 gp140 (Figure 3) was developed based on the process established for BG505 SOSIP.664 (Dey et al., 2018). Key activities included: (1) Lab-scale feasibility runs, (2) Evaluating the robustness of 2G12 (Polymun Scientific, Austria) capture chromatography, MabSelect SuRe (Cytiva, Marlborough, MA) affinity chromatography, and Capto adhere (Cytiva, Marlborough, MA) chromatography, and (3) Assessing harvest and process intermediate hold-time stability. The process was thoroughly evaluated for step yields, overall yield, and impurity removal, including both product-related impurities (e.g., high molecular weight [HMW] and low molecular weight [LMW] species) and process-related impurities (e.g., residual host cell proteins [HCP], 2G12 ligand, and residual DNA). The final downstream process is described in detail in section 2.5.4.

**Figure 3:**
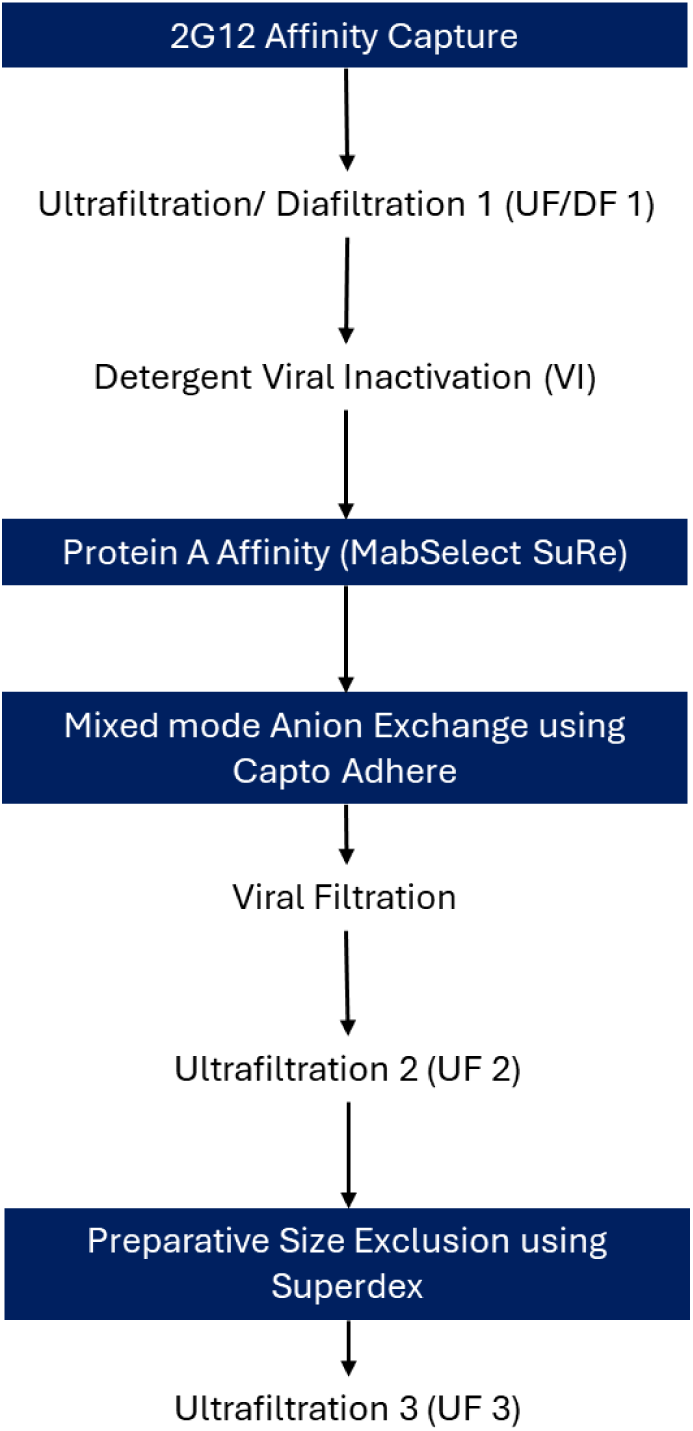
Feasibility Run – Downstream Process Flow Chart

#### 2.5.1 Process evaluation during Feasibility runs

After an initial process assessment, several parameters were identified to be modified for robust scale up. The full downstream process evaluated for the Feasibility Run is shown in Figure 3. During Feasibility Runs the MabSelect SuRe and Capto adhere chromatography steps were modified to be operated using a 20 cm bed height to ensure scalability of those unit operations. Additionally, to evaluate robustness all chromatography steps were executed with multiple cycles and using conditions that would encompass theoretical worst-case yield or worst-case product quality. Parameters such as column loading, pH, conductivity and elution buffer concentrations were considered within the design as listed in Table 3 for testing process robustness.

**Table 3:** Robustness Evaluations and Operational Conditions.

| Chromatography Unit Operation | Robustness Study | Operational Parameter 1 | Operational Parameter 2 | Operational Parameter 3 |
| --- | --- | --- | --- | --- |
| 2G12<br>(affinity capture) | Theoretical WCPQ | Elution Buffer<br>3.2 M MgCl<sub>2</sub> | High Loading<br>1.3 g/L<sub>resin</sub> | N/A |
|  | Theoretical WCY | Elution Buffer<br>2.8 M MgCl<sub>2</sub> | Low Loading<br>0.75 g/L<sub>resin</sub> | N/A |
| MabSelect SuRe (flowthrough) | Theoretical WCPQ | Not tested in this run | Not tested in this run | Not tested in this run |
|  | Theoretical WCY | Low Loading<br>5.0 g/L<sub>resin</sub> | N/A | N/A |
| Capto adhere (flowthrough) | Theoretical WCPQ | pH 7.2 | 13.0 mS/cm | High Loading<br>5.0 g/L<sub>resin</sub> |
|  | Theoretical WCY | pH 7.4 | 9.0 mS/cm | Low Loading<br>2.0 g/L<sub>resin</sub> |
| Superdex 200pg | Theoretical WCY | Low Loading<br>1.0% CV | N/A | N/A |

Lastly, the opportunity was utilized to evaluate Planova 20N (Asahi Kasei, MI, USA) virus filtration using fresh and freeze/thawed load material to enable any future viral clearance and scale down studies after the cGMP batch was completed.

#### 2.5.2 Evaluation of Removal of Preparative SEC chromatography

Due to supply constraints of the Superdex (Cytiva, Marlborough, MA) 200pg size exclusion resin (preparative SEC), it was pertinent to the cGMP campaign timeline to move forward without this unit operation. A small-scale study was performed using Demonstration Run (Pilot-scale) preparative SEC load material to evaluate whether the unit operation could be removed from the process without negatively impacting product quality. Key considerations were residual HCP clearance, SEC-HPLC % LMW species, and RP-HPLC % pre-Main species.

#### 2.5.3 Downstream Process Intermediate Hold Time Stability Study

During the pilot-scale Demonstration Run, the downstream process intermediate hold time stability was assessed to establish acceptable hold times for each intermediate at both ambient temperature (15 – 25°C) and 2 – 8°C to support cGMP manufacturing. Table 4 shows the process intermediates tested for hold time stability and any equivalent intermediates in the downstream process. The process intermediates which had the same buffer matrix, pH, and similar protein concentrations were considered equivalent for stability purposes. For the purposes of supporting manufacturing, SE-HPLC and relative potency were the two analytical outputs assessed.

**Table 4:**
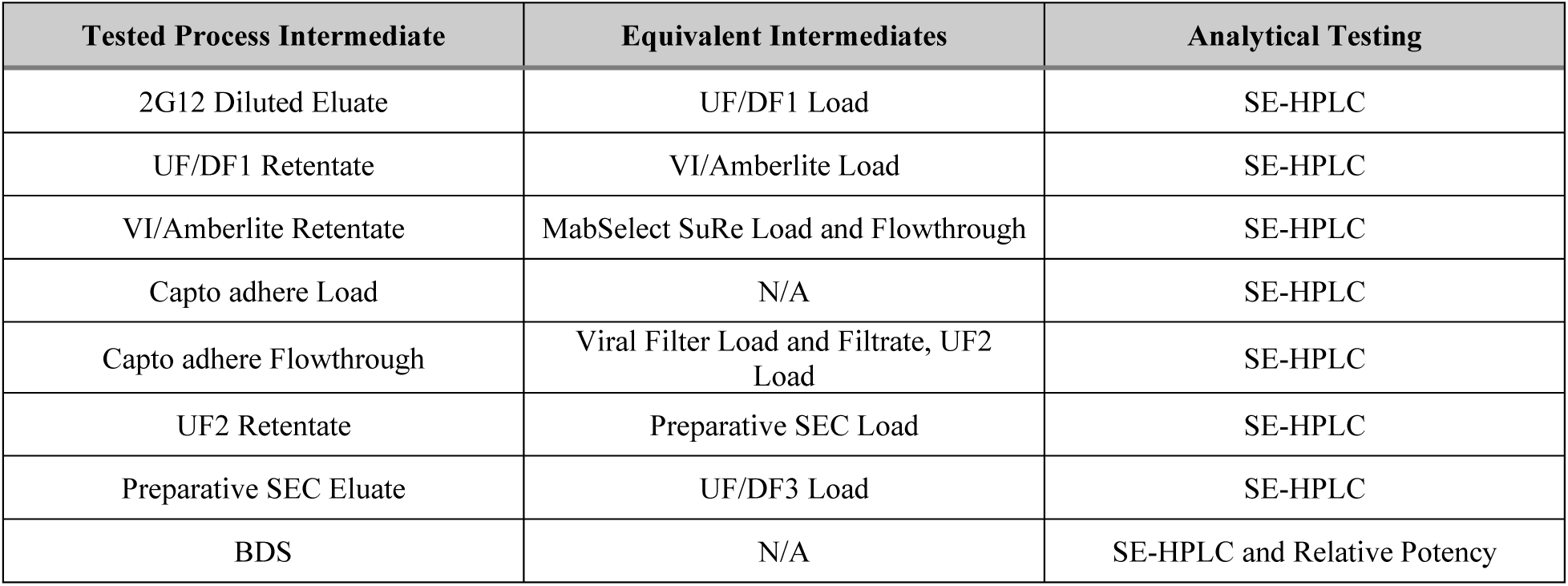
Process Intermediate Hold Time Stability Study.

SE-HPLC analysis was performed for all intermediates listed in Table 4. To establish hold time criteria, an intermediate was considered unstable if the % HMW or % LMW increased by more than 0.7%. Relative potency was performed for the initial and final timepoints for each temperature of the BDS. To establish hold time data, the BDS was considered unstable if the relative potency varied by more than 15% over the assessed hold time.

#### 2.5.4 The final process for pilot scale and GMP scale

After downstream process development and robustness evaluations, the process was finalized and transferred for the cGMP manufacturing (Figure 4). The final process had following unit operations.

**Figure 4:**
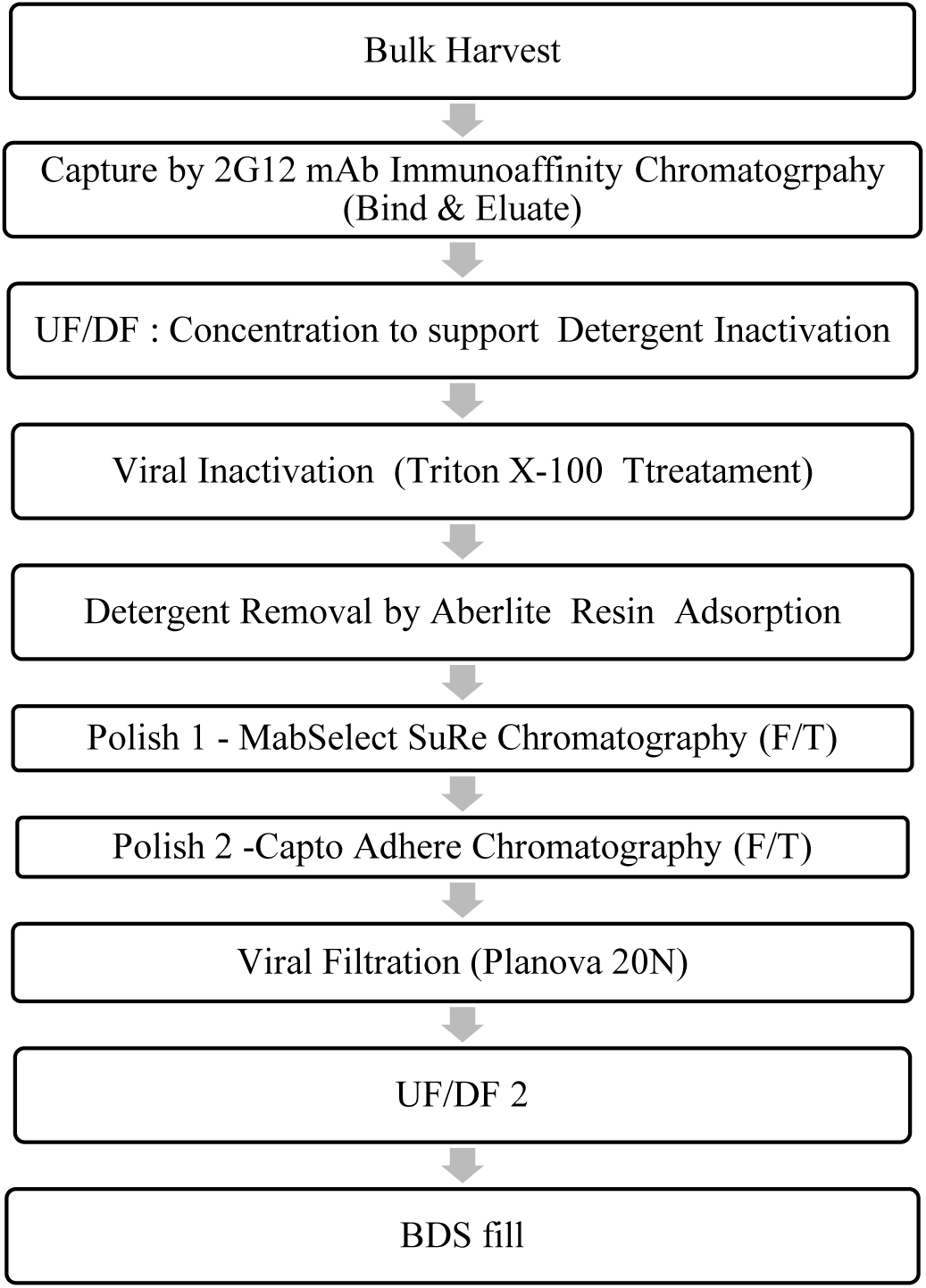
Purification Process for Manufacturing N332-GT5 gp140 Drug Substance.

**Figure 5:**
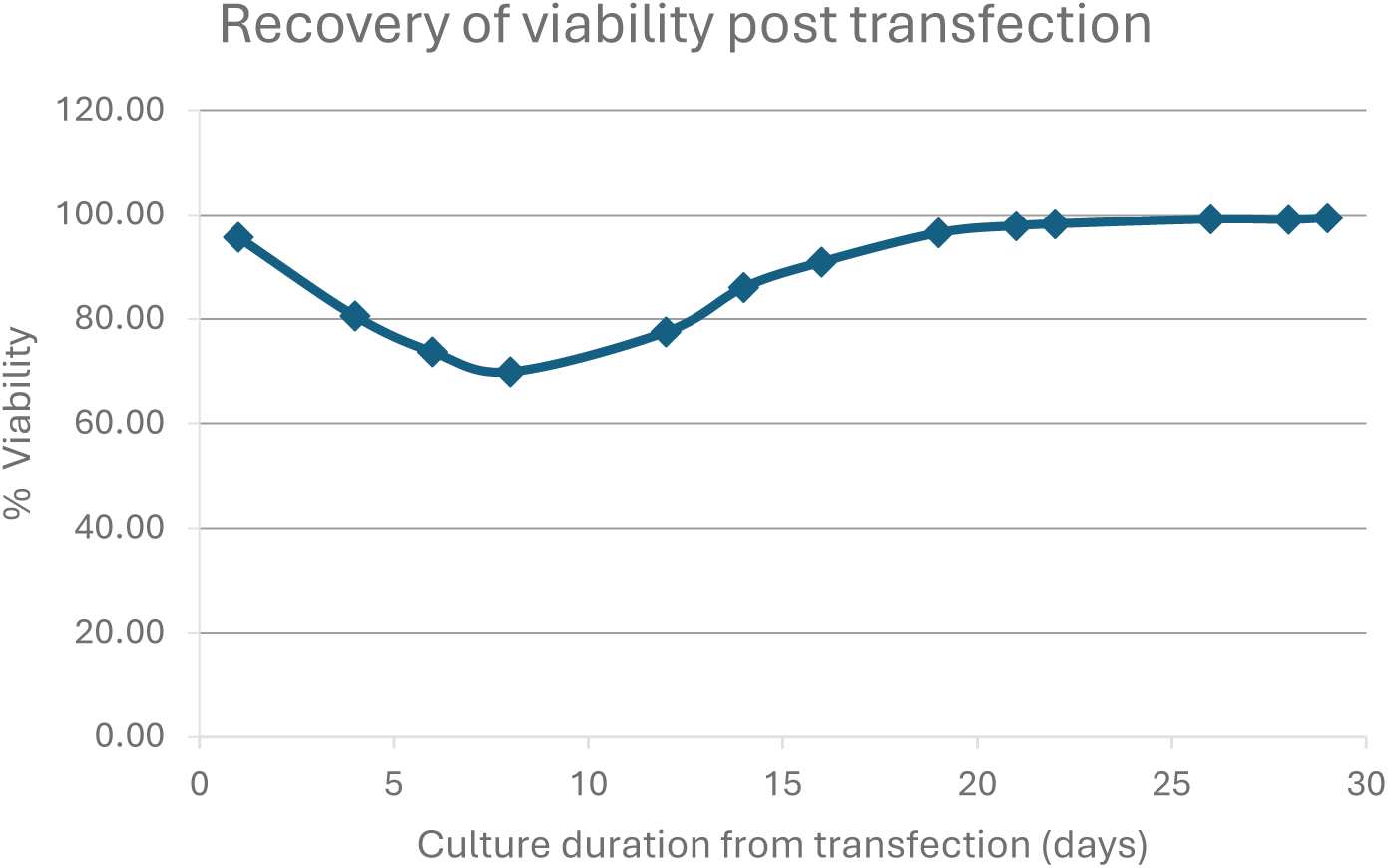
Recovery of the glutamine-free media selected stable pool expressing N332-GT5 gp140

##### 2.5.4.1 2G12 Chromatography

The clarified harvest was loaded onto a 2G12 affinity chromatography column, pre-equilibrated with 20 mM acetate buffer containing 2% v/v benzyl alcohol (pH 5.0). Before use, the column was sanitized, held for 12–16 h, and then equilibrated. After loading, the column was washed with phosphate-buffered saline (PBS, pH 7.4) to remove impurities, and the product was eluted with 50 mM Tris and 3 M MgCl₂ (pH 7.2). The eluate was diluted directly into 20 mM Tris, 75 mM NaCl (pH 8.0) and shown to be stable for up to 3 days at 15–25 °C and 7 days at 2–8 °C.

##### 2.5.4.2 UF/DF #1

The 2G12 eluate was concentrated and buffer-exchanged using a Pellicon Ultracel 100 kDa C-screen membrane (MilliporeSigma, MA, USA). The cassette was sanitized with 0.5 M NaOH for 30–60 min, flushed with WFI, and equilibrated in 20 mM Tris, 75 mM NaCl (pH 8.0). The product was loaded at ≤ 50 g/m², concentrated to 2.3 ± 0.2 g/L, and diafiltered against the same buffer for at least six diavolumes (DV) at a TMP of 10–15 psi and a feed flow rate of 5–7 L/m²/min. The final retentate was adjusted to ∼2.0 g/L and remained stable for 1 day at 15–25 °C and 1 day at 2–8 °C.

##### 2.5.4.3 Detergent Viral Inactivation

Buffer-exchanged material was treated with 0.5% (v/v) Triton X-100 (SAFC, St. Louis, MO) prepared from a 10% (v/v) stock solution to inactivate potential viral contaminants. The solution was mixed for 5 min and held statically for 60–90 min at 20 ± 5 °C (pH 8.1). The inactivation was quenched by loading onto a pre-equilibrated Amberlite adsorption resin.

##### 2.5.4.4 Amberlite Resin Adsorption

Amberlite XAD-2 (MilliporeSigma, MA, USA) resin was used to adsorb and remove Triton X-100 detergent from the product pool. The resin was sanitized with 0.5 M NaOH and equilibrated with 20 mM Tris, 75 mM NaCl (pH 8.0) before use. The inactivated material was circulated over the resin for 120–150 min at room temperature. The effluent was collected, yielding a clear, colorless solution (pH=8.1; conductivity = 8.1 mS/cm) that remained stable for up to 2 days at 15–25 °C and for 7 days at 2–8 °C.

##### 2.5.4.5 MabSelect SuRe Chromatography

The detergent-removed intermediate was further purified using MabSelect SuRe Protein A resin operated in flow-through mode. The column was sanitized with 0.1 M NaOH, equilibrated in 20 mM Tris, 75 mM NaCl (pH 8.0), and the load passed through at 20 °C. Product collection was defined between ≥ 0.05 AU/cm and ≤ 0.05 AU/cm, including collection of the flow-through and a portion of the wash step.

##### 2.5.4.6 Capto adhere Chromatography

The Protein A flow-through was polished by flow-through chromatography using Capto adhere resin. The column was sanitized with 0.5 M NaOH, equilibrated with 110 mM Tris, 75 mM NaCl (pH 7.2), and loaded with material adjusted to pH 7.30 ± 0.10 and conductivity 11.0 ± 0.2 mS/cm. Product collection was performed between ≥ 0.05 AU/cm and ≤ 0.15 AU/cm. The intermediate was stable for 3 days at 15–25 °C and 7 days at 2–8 °C.

##### 2.5.4.7 Virus Filtration

Virus filtration was performed using a Planova 20N virus filter (Asahi Kasei) with a 0.1 µm Sartopore 2 prefilter (Sartorius, Gottingen, Germany). Filters were flushed and equilibrated with 110 mM Tris, 75 mM NaCl (pH 7.2) prior to use. Filtration was performed at 10–12 psi, followed by a buffer flush to recover residual product. The process step achieved a 98% product yield, and both load and filtrate were stable for 3 days at 15–25 °C and 7 days at 2–8 °C.

##### 2.5.4.8 UF/DF #2

A final ultrafiltration/diafiltration step using a Pellicon Ultracel 100 kDa C-screen membrane (MilliporeSigma, MA, USA) was used for buffer exchange into 20 mM Tris, 100 mM NaCl (pH 7.5). The cassette was sanitized with 0.5 M NaOH (30–60 min), flushed with WFI, and equilibrated with diafiltration buffer. The viral filtrate was loaded at ≤ 100 g/m², concentrated to 2.0 g/L, diafiltered for ≥ 10 DV, and diluted to 1.0 g/L. The final product was stable for 3 days at 15–25 °C and 7 days at 2–8 °C.

### 2.6 Viral Clearance Studies

To ensure viral clearance, four specific downstream unit operations were included in the purification process of N332-GT-5 gp140. These steps involved: (i) 2G12 affinity column chromatography, (ii) Triton X-100 detergent inactivation, (iii) Capto adhere column chromatography, and (iv) Nanofiltration using Planova 20N. A scale-down model of the purification process was developed to ensure comparability with the manufacturing-scale process. To assess the effectiveness of these steps in clearing viruses, a viral clearance study was conducted using material from the cGMP run and included two model viruses—Xenotropic Murine Leukemia Virus (XMuLV), an enveloped retrovirus, and Minute Virus of Mice (MMV), a non-enveloped adventitious virus. Each unit operation was performed separately, with known quantities of virus spiked into representative load materials. Viral clearance was calculated by measuring the log (base 10) reduction of virus (LRV) for each unit operation, and these individual clearances were summed to determine the total viral reduction across the process. The virus reduction data provides an estimate of the potential risk of viral contamination in the final product.

### 2.7 Product Characterization

#### 2.7.1 Site-specific *N*-glycan Analysis

Two methods were used for site specific analysis of glycans. One method is DeGlyPHER, which generates peptides with mass signatures that code for the glycosylation status of the glycosite. The other method is a more traditional LC-MS glycoproteomic method that generates peptides with intact glycans, providing additional information on the glycoforms at each glycosite. Each of the two methods is briefly described below:

DeGlyPHER (Baboo et al., 2023) was used to determine site-specific glycan occupancy and processivity on the N332-GT5 trimer. Briefly, disulfide bonds on the glycoprotein were reduced and alkylated before digestion with Proteinase K, followed by sequential deglycosylation of glycopeptides with Endo H and then PNGase F in the presence of H_2_^18^O. The peptides were separated on C18 resin using an EASY-nLC 1200 UHPLC (Thermo) and nanosprayed into a Q Exactive HF-X mass spectrometer (Thermo). Spectra were acquired in data-dependent mode with HCD fragmentation. On the Integrated Proteomics Pipeline (IP2, Bruker), extracted tandem mass spectra were searched using ProLuCID (Xu et al., 2015) against the known protein sequence of N332-GT5 within a CHO (Chinese Hamster Ovary) cell proteome background, with C+57.02146 Da as a static modification and N+2.988261 Da (signature for complex glycans), N+203.079373 Da (signature for high-mannose/hybrid glycans), M+15.994915 Da, and N-terminal Q–17.026549 Da as variable modifications. These were filtered at the spectrum level to up to 1% FDR (Peng et al., 2003) using DTASelect2 (Tabb et al., 2002) and then quantified with Census2 (Park et al., 2008) label-free analysis, applying “match between runs.” GlycoMSQuant (Baboo et al., 2021) was used to compile the final results, aligning PNGS to Env of the HXB2 HIV-1 variant.

Traditional liquid chromatography (LC-MS) glycoproteomics analysis was performed using a Thermo Orbitrap Eclipse. Peptides and glycopeptides were generated from the provided glycoprotein using three proteases – trypsin, chymotrypsin, and alpha-lytic protease (Watanabe et al., 2020). For each replicate, data from LC-MS analyses of glycopeptide and peptide pools obtained with each protease were combined. Data was analyzed using Byos (Protein Metrics).

#### 2.7.2 Negative-stain electron microscopy (nsEM)

N332-GT5 from the 50L demonstration run was diluted in Tris-buffered saline (50 mM Tris pH 7.4, 150 mM NaCl) to a final concentration of 0.02 mg/mL and adsorbed onto glow-discharged, carbon-coated copper grids (Electron Microscopy Sciences). Excess solution was blotted with Whatman #1 filter paper and the grid was stained with 2% (w/v) uranyl formate for 45 s. Excess stain solution was blotted and 82 micrographs were collected on an FEI Tecnai Spirit TEM equipped with an FEI Eagle 4K CCD (120 kEv, 2.06 Å pixel size, 52,000 nominal magnification). Data collection was automated using Leginon (Suloway, C et al., 2005). Micrographs were imported into CryoSPARC (Pujani, et al., 2017), Blob Picker (minimum circular diameter 180 Å) was performed to locate particles, which were subsequently extracted at 160 pixel box size. This particle stack was subjected to two rounds of 2D classification, with a total of 6,086 particles analyzed for native-like trimeric properties by comparison to previously published HIV Env SOSIP production runs (Dey et al., 2018; Bale et al., 2025).

## 3 Results and Discussion

### 3.1 Cell Line Development and Characterization

#### 3.1.1 Stable Pool Development and Characterization

The recovery of the N332-transfected stable pool presents the typical Leap In transposase-mediated profile characterized by a relatively low viability drop and fast recovery.

A N332-GT5 stable pool was used to initiate a 14-day fed-batch culture. The viability and VCD values during the run are shown in Figure 6. The productivity and the product quality were assessed by PGT145 and BG18 Octet measurements (Figure 7) and reduced SDS PAGE analysis (Figure 8).

**Figure 6:**
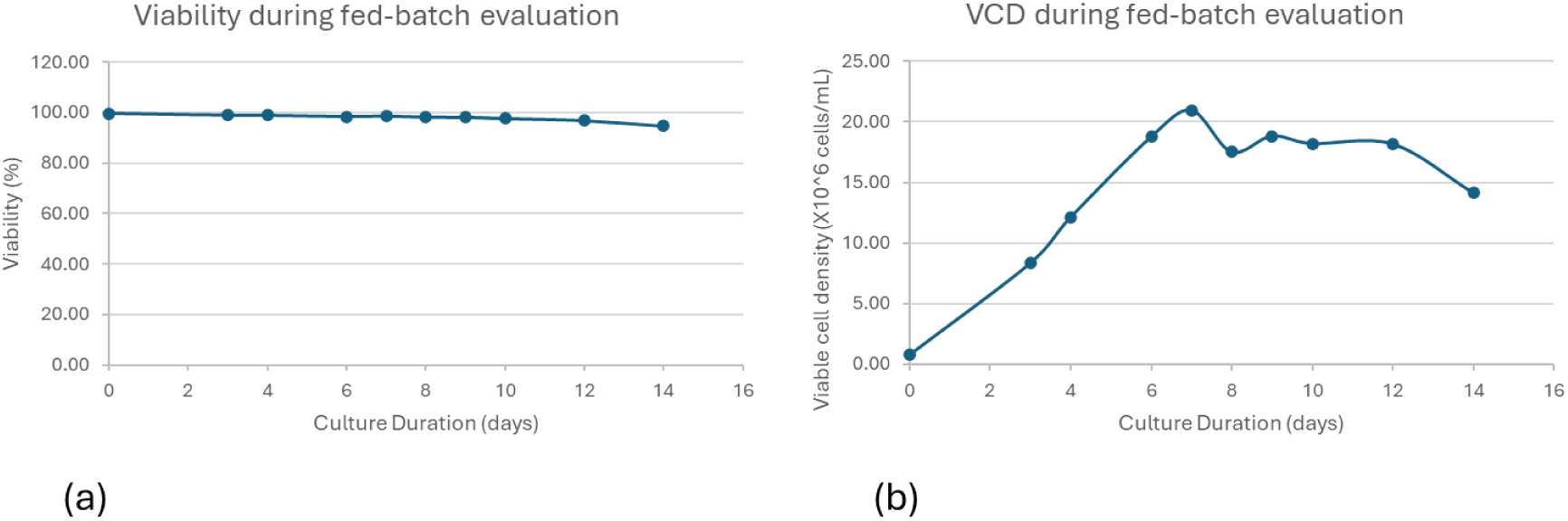
The viability (a) and VCD (b) profiles during the 14-day fed batch of the N322 stable pool. The graphs show healthy, robust cell growth with no signs of potential significant product-related growth inhibition.

**Figure 7:**
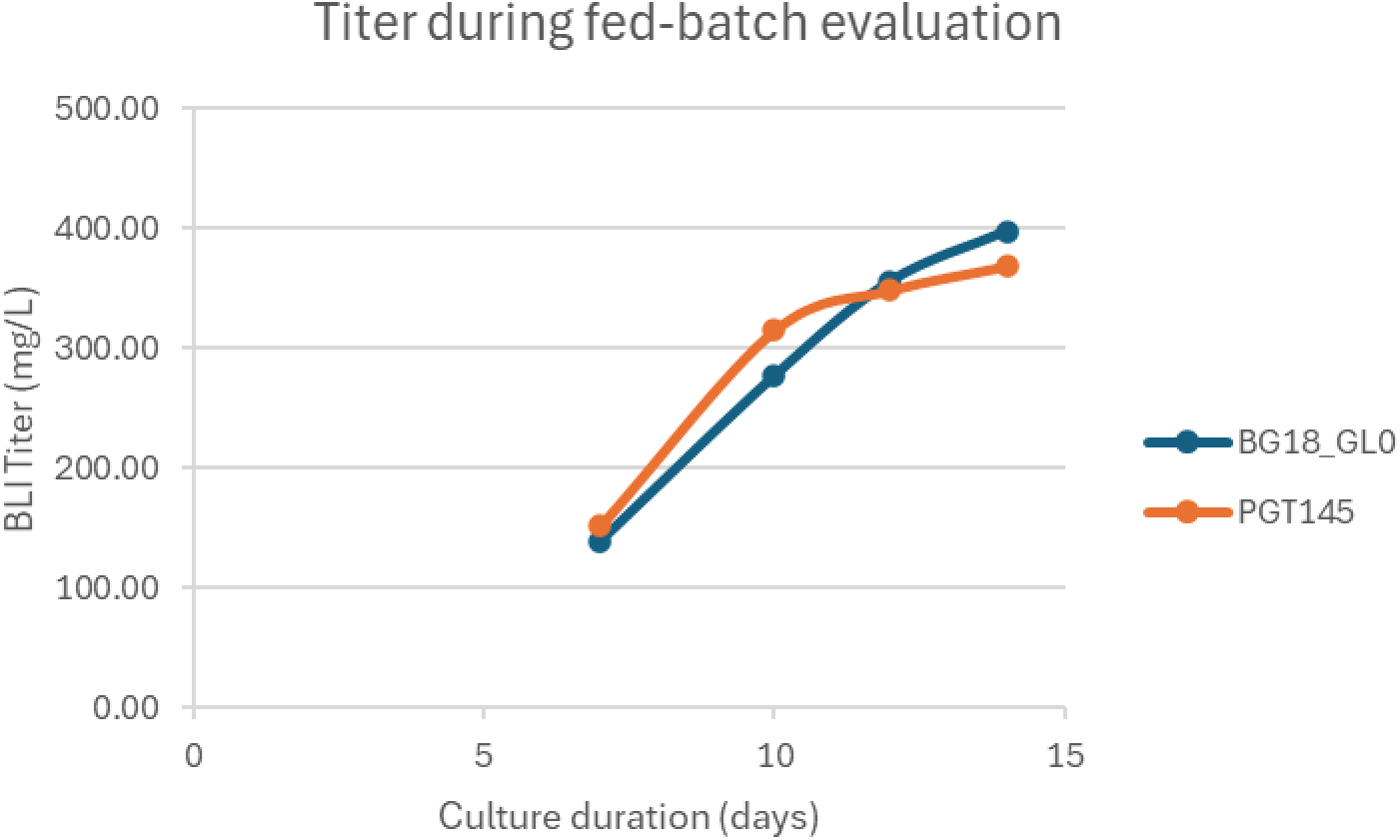
BLI titer measurement

**Figure 8:**
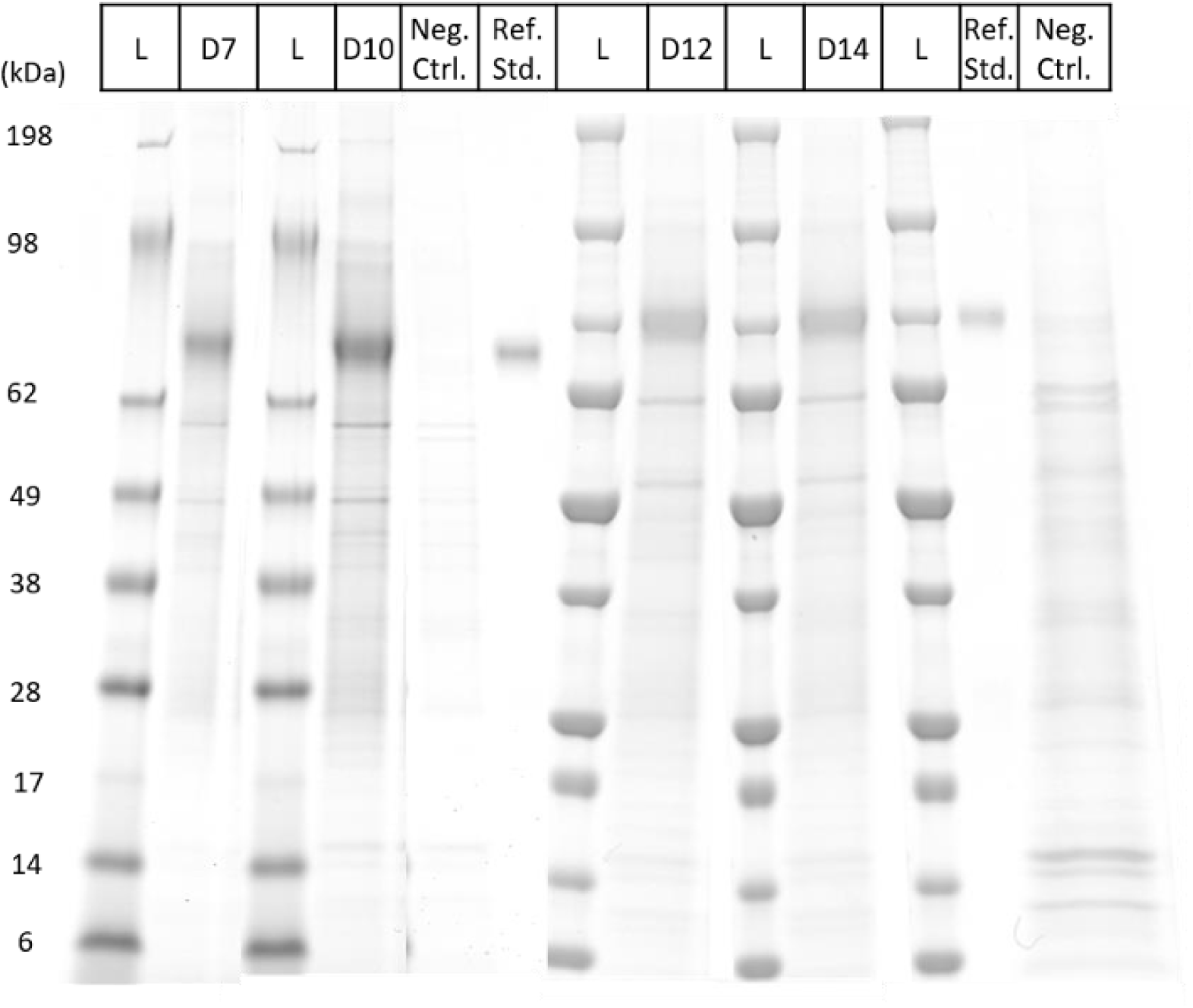
Reduced SDS PAGE for product quality assessment: The reduced SDS PAGE patterns in the clarified harvests are comparable to the purified reference standard. There is no indication of significant proteolysis, and the furin cleavage seems close to complete as well.

To evaluate the productivity and quality of the recombinant protein produced by the N332 stable pool, samples were taken from the fed-batch culture on days 7, 10, and 12, and the culture was harvested on day 14. Detectable product levels of BG18 and PGT145 (Figure 7) were observed, along with reduced SDS-PAGE analysis (Figure 8), in the clarified harvests collected. Figure 7 shows the comparable detectable productivities of BG18 and PGT145 indicating that the main form of N332-GT5 gp140 in the harvest material is the intended trimer. Additionally, considering the product’s complexity, a productivity of 350-400 mg/L was deemed acceptable. The SDS-PAGE gels were run according to the methods section, with 3.25 µL of supernatant loaded into each lane for days 7 and 10, and 2 µL for days 12 and 14.

The comparable BG18 and PGT145 detectable productivities indicate that in the harvest material, the predominant form of N332-GT5 gp140 is the intended trimer. Also, given the product’s complexity, a productivity of 350-400 mg/L was acceptable.

The SDS-PAGE gels were run as described in the methods section, where 3.25 µL of supernatant was loaded into each lane for D7 and D10 samples, and 2 µL of supernatant was loaded for D12 and D14 samples.

#### 3.1.2 First Clone Ranking

213 different clones were ranked by detectable productivity in seven-day fed-batch cultures using the BG18 method. The end-point viability of most clones exceeded 80%. BG18_GL0 detectable productivity was measured in the Day 7 clarified harvests (Figure 9a). Using their BG18_GL0 titer, the top 50 clones, along with three clones from the middle productivity range and three from the lower range, were selected for the PGT145-based Octet/BLI assay to determine titer (Figure 9b).

**Figure 9:**
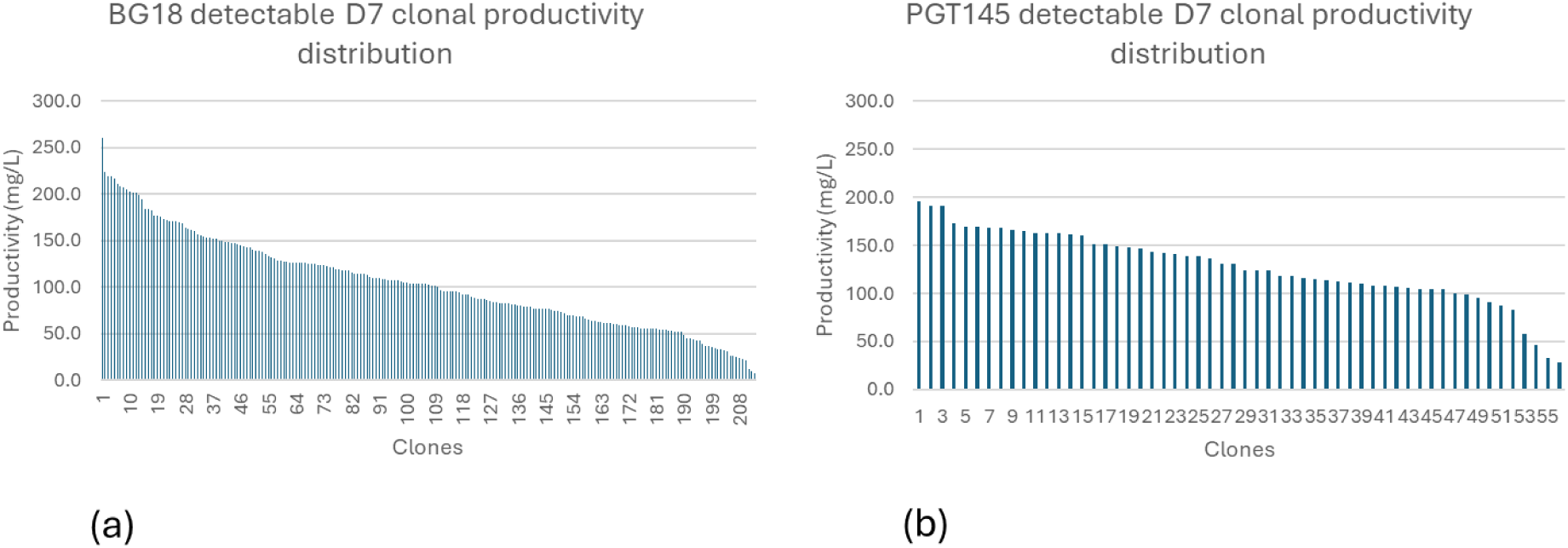
PGT145-based Octet/BLI assay

The clonal productivity distributions shown in Figure 9 illustrate typical Leap-In mediated profiles. Based on their growth patterns and BG18/PGT145 productivity ratios, 24 clones were chosen for a second round of clone ranking. The ranking criteria for these 24 clones are listed in Table 5.

**Table 5:** Ranking parameters for the top 24 clones selected for further analysis.

| Clone ID | Day7-Viability[%] | Day7-Viable Cell Count[e6cells/ml] | Day7-BG18_GL0 Titer[mg/L] | Day7-PGT145 Titer[mg/L] | Ratio PGT145/BG18_GL0 |
| --- | --- | --- | --- | --- | --- |
| C311 | 97.85 | 23.30 | 260.6 | 195.9 | 0.8 |
| C1051 | 96.58 | 19.04 | **216.4** | 191.5 | 0.9 |
| C524 | 98.60 | 22.17 | 198.9 | 191.3 | 1.0 |
| C261 | 98.06 | 22.23 | 194.1 | 173.4 | 0.9 |
| C845 | 96.03 | 19.70 | 201.1 | 169.4 | 0.8 |
| C273 | 95.33 | 10.46 | 218.7 | 169.3 | 0.8 |
| C464 | 95.76 | 17.29 | 219.6 | 168.5 | 0.8 |
| C1437 | 99.31 | 28.60 | 149.6 | 168.2 | 1.1 |
| C484 | 97.23 | 21.79 | 176.1 | 166.3 | 0.9 |
| C404 | 96.93 | 18.78 | 156.4 | 164.6 | 1.1 |
| C100 | 98.48 | 18.30 | 168.1 | 162.8 | 1.0 |
| C171 | 96.62 | 14.99 | 148.3 | 162.6 | 1.1 |
| C1436 | 99.52 | 24.02 | 150.2 | 162.1 | 1.1 |
| C134 | 97.01 | 19.48 | 207.6 | 161.8 | 0.8 |
| C1099 | 99.37 | 23.57 | 172.5 | 160.8 | 0.9 |
| C371 | 99.13 | 22.64 | 177.0 | 151.7 | 0.9 |
| C280 | 97.74 | 23.15 | 205.3 | 151.6 | 0.7 |
| C167 | 96.88 | 18.31 | 208.3 | 147.9 | 0.7 |
| C235 | 96.66 | 19.23 | 223.7 | 142.3 | 0.6 |
| C701 | 98.42 | 13.17 | 156.0 | 136.7 | 0.9 |
| C1183 | 98.07 | 14.06 | 153.2 | 124.0 | 0.8 |
| C584 | 98.29 | 22.74 | 142.3 | 118.1 | 0.8 |
| C1309 | 99.07 | 18.69 | 140.2 | 115.4 | 0.8 |
| C29 | 99.33 | 15.91 | 102.1 | 99.6 | 1.0 |

#### 3.1.3 Second Clone Ranking

The second clone ranking step for the selected 24 clones was performed using a 14-day fed-batch process in TubeSpins at a 10 mL volume. The performance of clones was assessed by growth, viability, titer by binding assay, and product quality assessment by SDS-PAGE gels. Peak VCD ranged from 15 to 25 × 10^6^ cells/mL for all clones. Cultures were harvested when the measured viability was greater than 80%.

Figure 10 presents the detectable productivities of BG18 and PGT145 during the second clone ranking.

**Figure 10:**
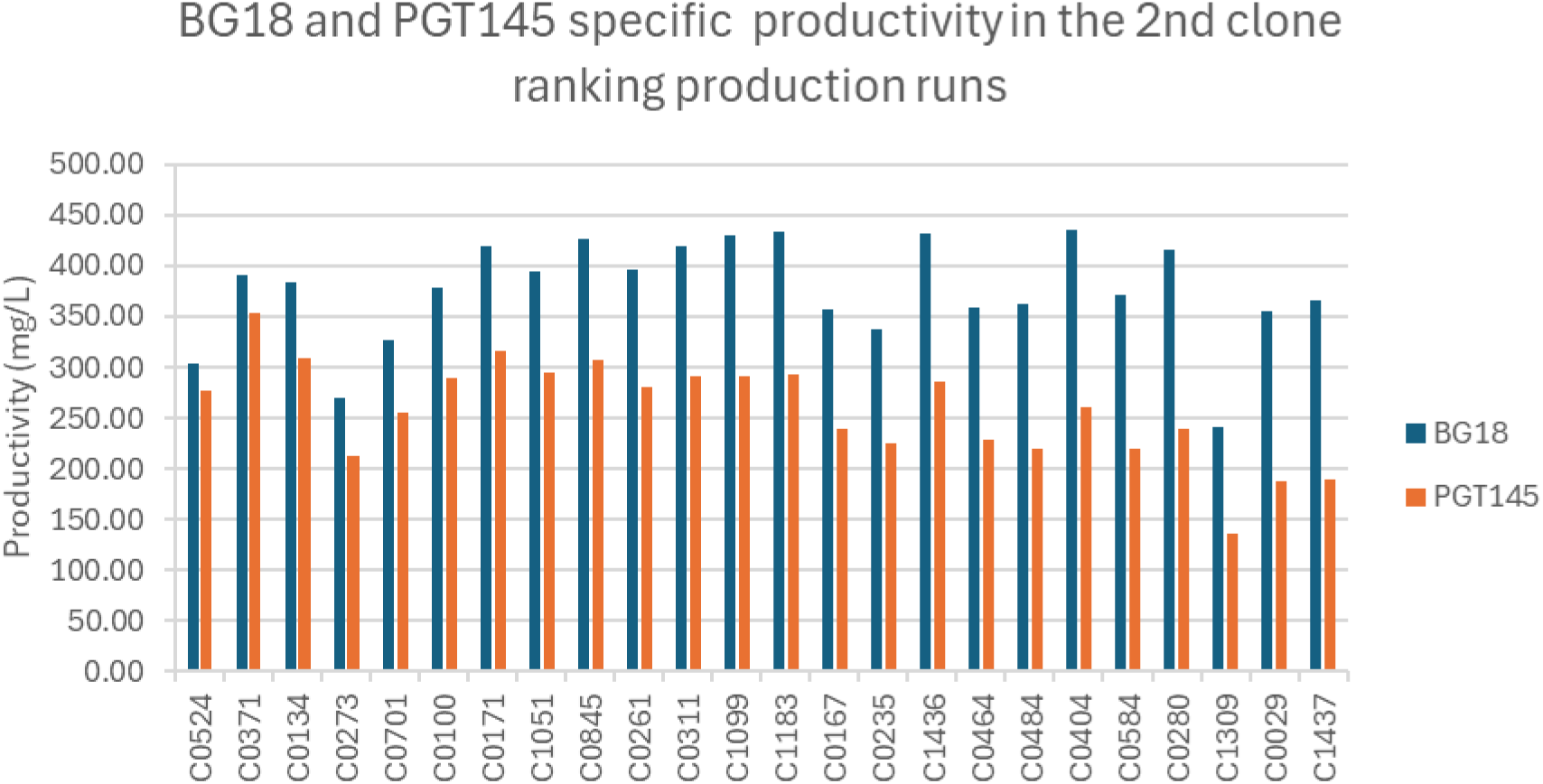
The BG18 and PGT145 detectable productivities in the top 24 clones during the second clone ranking fed batch run.

Product quality was also assessed by reduced SDS-PAGE analysis, which monitored furin cleavage in the individual clones. Presented in Figure 11.

**Figure 11:**
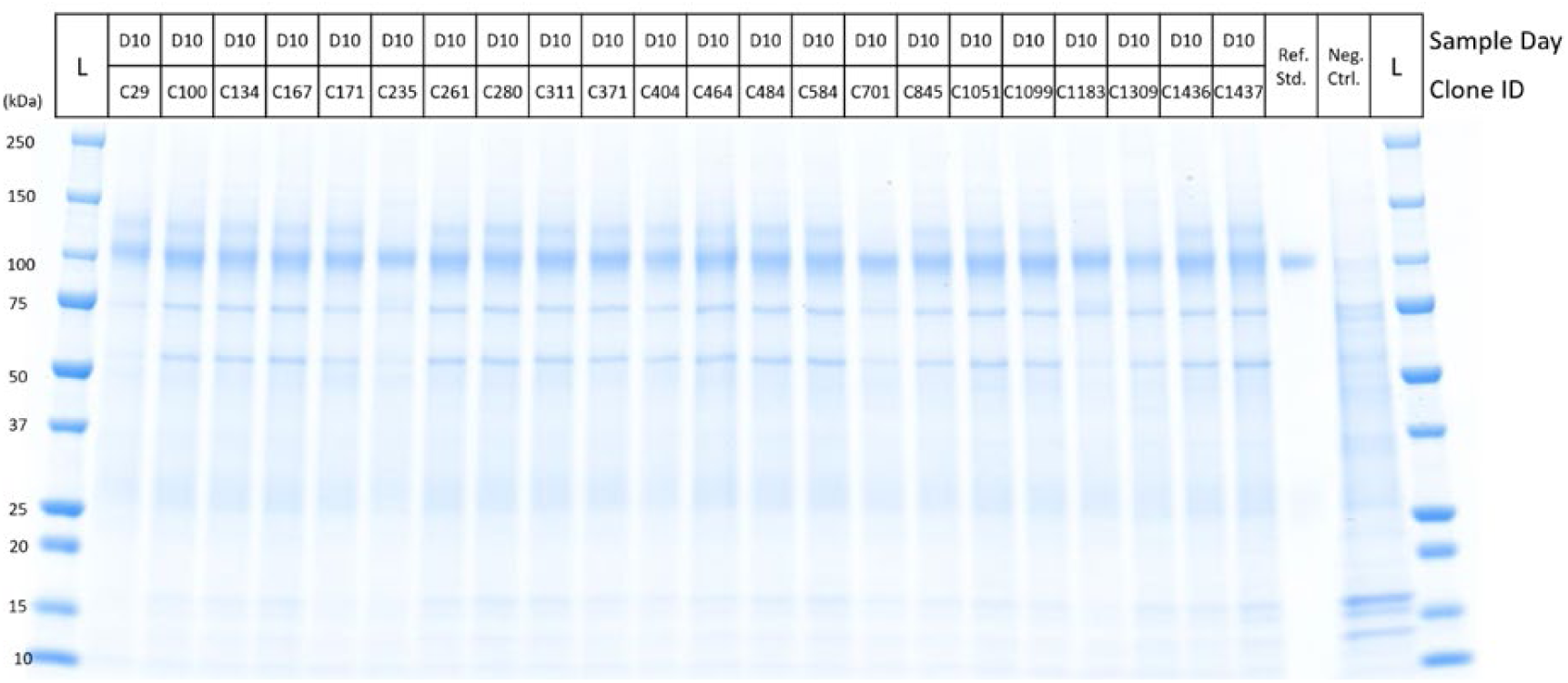
Reduced SDS PAGE analysis of the 24 selected clones.

The absence of the faint band migrating above the furin cleaved expected product indicates complete furin cleavage. Based on growth characteristics, productivity, product quality and the presence of recombinant furin, clone C235 has been selected for upstream process development.

### 3.2 Cell Culture Process Establishment

#### 3.2.1 Direct Medium Adaptation Performance

The performance of clone C235 during direct adaptation from EX-CELL AFB to Dynamis medium was evaluated by monitoring doubling time and cell viability across five passages.

As shown in Figure 12 The doubling time after thawing in EX-CELL AFB medium was 23.1 hours, which decreased to 21.2 hours at passage 2 following transfer into Dynamis medium. Doubling times gradually increased at subsequent passages, reaching 25.9, 27.0, and 28.7 hours for passages 3, 4, and 5, respectively. The modest increase in doubling time at later passages likely reflects continued physiological adjustment to the new medium composition. Cell viability remained high throughout the adaptation process (Figure 12). Viability at thaw (passage 1, EX-CELL AFB) was 100% and remained above 98% through passage 4 in Dynamis medium (99%, 98%, and 98%, respectively). A slight reduction to 96% was observed at passage 5, which remained well within acceptable limits for process development.

**Figure 12:**
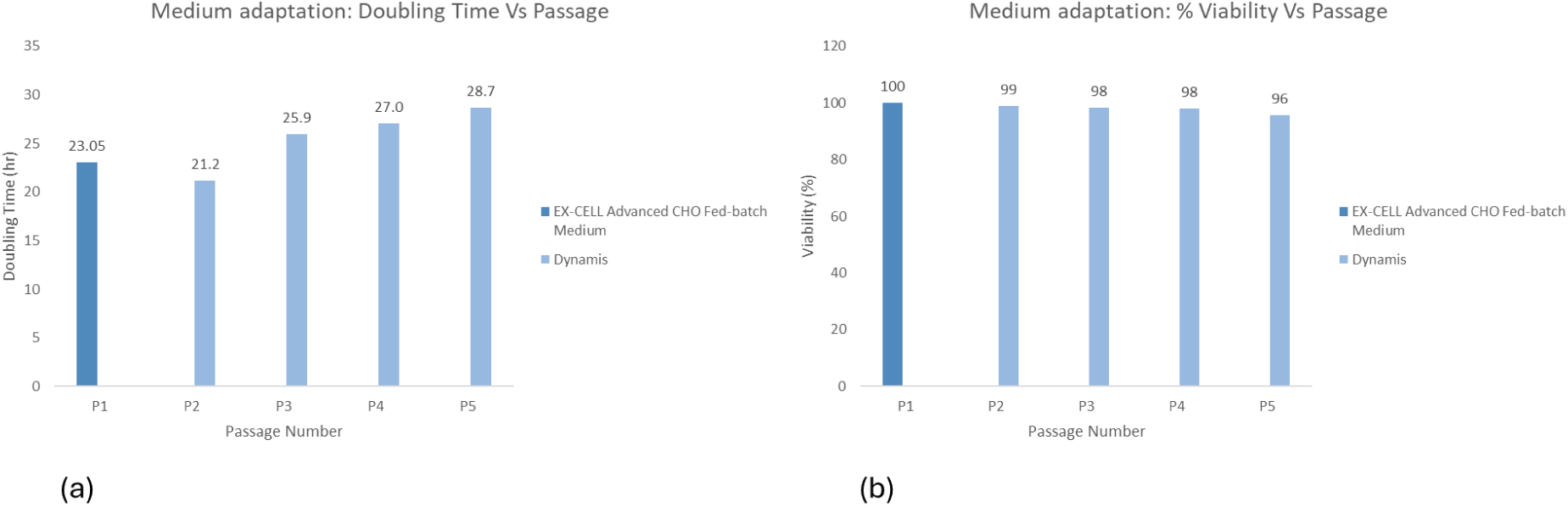
Medium adaptation cell culture data. (a) Doubling time (hours) for clone C235 during direct adaptation from EX-CELL AFB to Dynamis medium; (b) Cell viability (%) for clone C235 during direct adaptation from EX-CELL AFB to Dynamis medium.

Overall, clone C235 exhibited robust growth kinetics and sustained high viability during direct adaptation to Dynamis medium, confirming that the cell line is well-suited for propagation and process development in this commercially available, chemically defined formulation. Passage 5 was therefore used to generate Dynamis-based RCB material and initiate subsequent upstream process optimization.

#### 3.2.2 Ambr250 Process Optimization

Peak viable cell concentrations (VCC) and Day 14 VCC values are summarized in Table 6. All cultures exhibited rapid growth during the first 5–8 days, followed by a decline in viability after Day 10. The highest peak VCC (25.2 × 10^6^ cells/mL) was achieved in the *High inoc, 40% CB7a slow feed, 34 °C Day 5* condition on Day 6. In comparison, *Control #1* and *Control #2* reached peak VCCs of 23.0 × 10^6^ cells/mL on Days 8 and 7, respectively. The lowest peak VCC (16.5 × 10^6^ cells/mL on Day 9) was observed under the *pH 6.90* condition. By Day 14, the *34 °C Day 5* condition maintained the highest VCC (18.6 × 10^6^ cells/mL), indicating improved culture longevity relative to the controls (*Control #1*: 15.3 × 10^6^; *Control #2*: 15.5 × 10^6^ cells/mL). The *35% CB7a bolus* condition resulted in the lowest Day 14 VCC (11.0 × 10^6^ cells/mL), suggesting that higher bolus feed concentrations may have negatively impacted final cell viability.

**Table 6:** Cell Growth and % Viability values from Ambr^®^250 process optimization.

| Condition | Peak VCC | Peak VCC Day | Day 14 VCC | Day 14 Viability (%) |
| --- | --- | --- | --- | --- |
| *Control #1* | 23 | 8 | 15.3 | 81.7 |
| *Control #2* | 23 | 7 | 15.5 | 85.7 |
| *Control + 7a bolus* | 24 | 7 | 15.8 | 87.2 |
| *35% 7a slow* | 20.5 | 10 | 14.6 | 85.6 |
| *35% 7a bolus* | 21 | 7 | 11.0 | 78.5 |
| *High inoc, 40% 7a slow* | 22 | 5 | 12.8 | 79.6 |
| *High inoc, 40% 7a bolus* | 21.2 | 5 | 11.3 | 81.9 |
| *High inoc, 40% 7a slow, 34C Day 5* | 25.2 | 6 | 12.5 | 76.1 |
| *34C Day 5* | 24.6 | 7 | 18.6 | 85.9 |
| *pH 6.90* | 16.5 | 9 | 12 | 84.4 |
| *pH 6.90, pH 7.20 Day 6* | 18.7 | 8 | 12.4 | 76.2 |
| *pH 7.00, pH 7.20 Day 6* | 19.9 | 5 | 13.5 | 83 |

Collectively, these results indicate that an earlier temperature shift (Day 5) and higher initial inoculation density enhanced both peak and sustained cell growth. The *Control + 7a bolus* condition had the highest Day 14 viability of 87.2%.

The expression of N332-GT5gp140 was measured using biolayer interferometry (BLI) on an Octet system (Sartorius). The assay utilized the IAVI/Scripps N332-GT5 reference standard (1.0 mg/mL, Lot 19Apr0088). BLI titer profiles across the 12 ambr^®^250 bioreactors are shown in Figure 13. Among all tested conditions, the highest Day 14 titers were achieved with *Control + 7a bolus* (752 mg/L), *High inoculation with 40% 7a bolus* (721 mg/L), and *35% 7a bolus* (715 mg/L). The performance of the *Control + 7a bolus* condition surpassed both controls (*Control #1*: approximately 640 mg/L; *Control #2*: approximately 650 mg/L), indicating that supplementation with a single CB7a bolus feed enhanced overall productivity compared to the baseline feeding strategy. Based on these results, the *Control + 7a bolus* condition was selected for subsequent scale-up in the 50-L bioreactor run using Clone C235 RCB.

**Figure 13:**
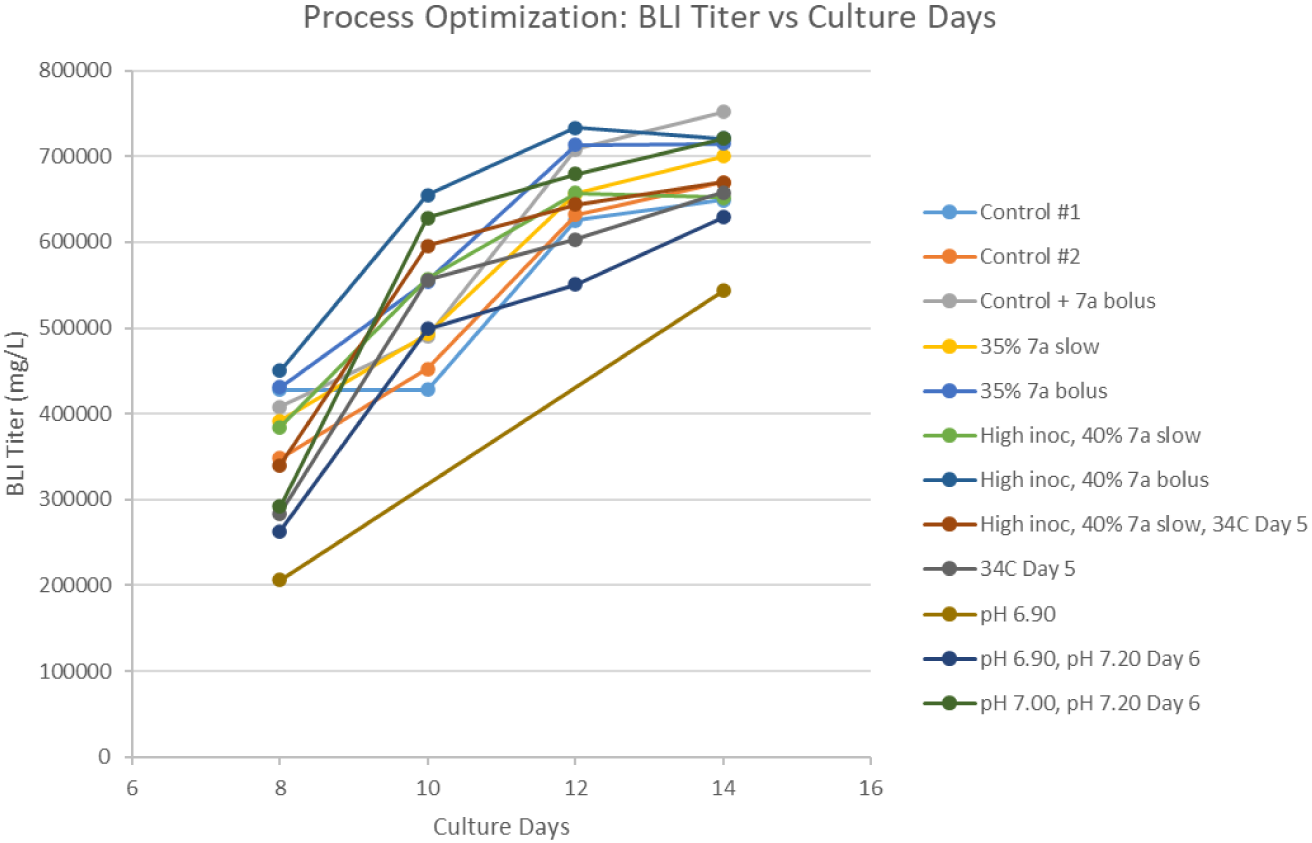
Ambr®250 Process Optimization BLI titer results (mg/L)

#### 3.2.3 Pilot and GMP scale runs

Two sequential 50-L fed-batch studies were performed using Clone C235 to evaluate the scalability and reproducibility of the N332-GT5-gp140 upstream process: a 50-L material supply run using the research cell bank (RCB) in an XDR-50, followed by a 50-L demonstration run using the master cell bank (MCB) in an XDR-200. Both runs used the *Control + 7a bolus* condition identified during process optimization. After the demonstration run, the same process was successfully executed for the GMP manufacturing using the XDR-200 bioreactor.

The XDR-50 (RCB) bioreactor was inoculated at 0.73 × 10^6^ cells/mL and reached a peak viable cell concentration (VCC) of 26 × 10^6^ cells/mL on Day 7. At harvest on Day 14, the culture had a VCC of 13.7 × 10^6^ cells/mL and a viability of 66.3%. In comparison, the XDR-200 (MCB) bioreactor was inoculated at 0.82 × 10^6^ cells/mL and achieved a peak VCC of 24.2 × 10^6^ cells/mL on Day 8, with a Day 14 VCC of 12.8 × 10^6^ cells/mL and a higher viability of 74.4%. For the GMP run, XDR-200 (MCB) bioreactor was inoculated at 0.76 × 10^6^ cells/mL and achieved a peak VCC of 20.33 × 10^6^ cells/mL on Day 8, with a Day 14 VCC of 13.3 × 10^6^ cells/mL and a viability of 77% (Figure 14). These data indicate that overall cell growth kinetics were comparable between the two scales and cell banks, with slightly improved end-point viability in the MCB run.

**Figure 14:**
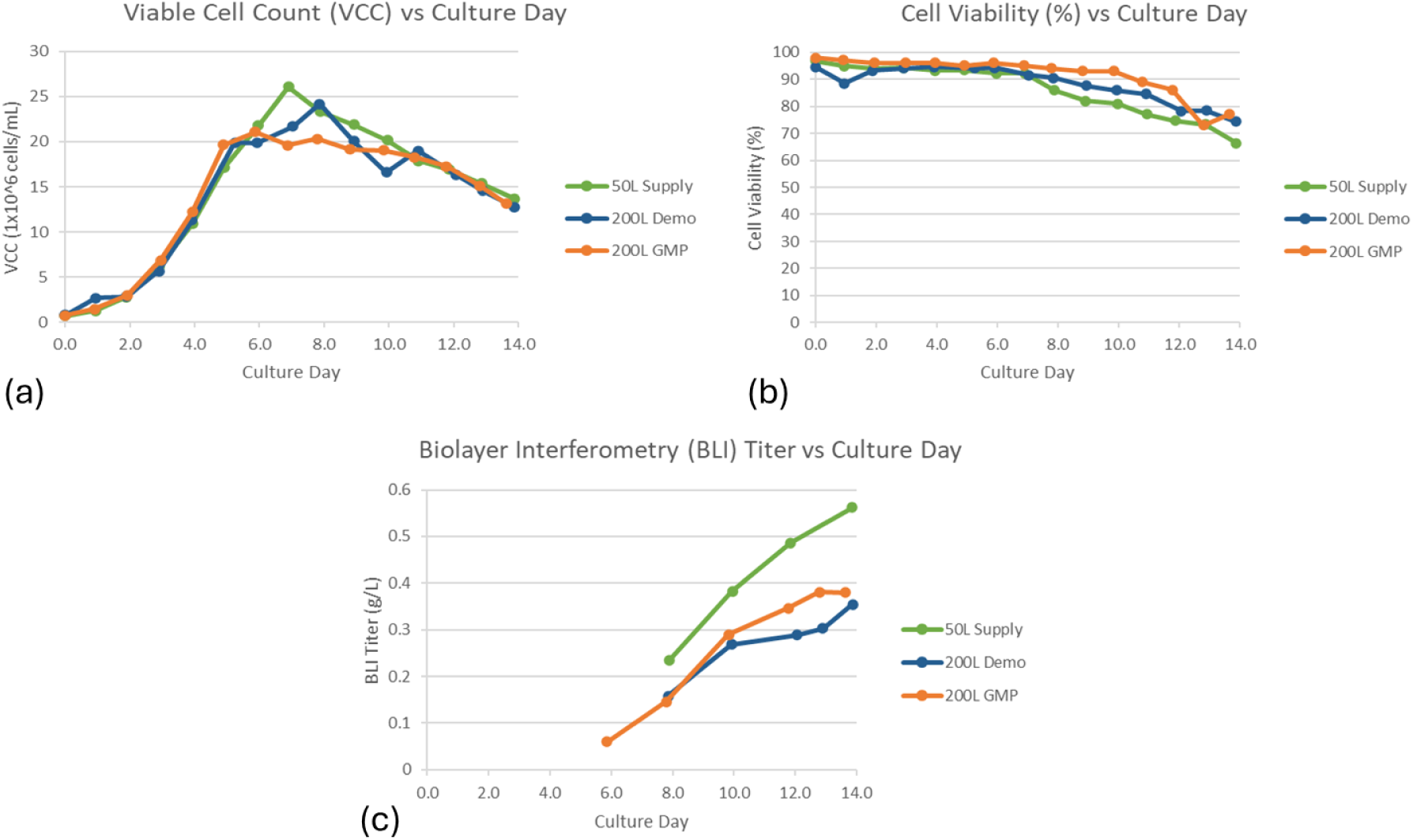
Cell culture performance during scale-up: (a)Viable Cell Count (VCC); (b) Cell Viability (%); (c) BLI titer for 50 L material supply run, 200L demonstration run and 200 L GMP run

Productivity was consistent with these growth and metabolic profiles (Figure 14). For the RCB run, the Day 14 bioreactor BLI titer was 562 mg/L, and approximately 57.7 kg of clarified harvest was recovered at 399 mg/L, corresponding to a harvest yield of 91.4%. For the MCB run, the Day 14 BLI titer in the bioreactor was 355 mg/L, and 69.0 kg of clarified harvest was obtained at 244 mg/L, yielding 90.8% harvest yield and a total N332-GT5 gp140 output of 16.8 g. For the GMP run using MCB, the Day 14 BLI titer in the bioreactor was 390.2 mg/L, and 87.6 kg of clarified harvest was obtained at 258 mg/L, yielding 91% harvest and a total N332-GT5 gp140 output of 22.6 g. The difference in titer was mainly due to the reference material used. The 50L Supply (C235 RCB) BLI titer was determined using IAVI/Scripps N332-GT5 reference material, 1.0 mg/mL, Lot: 19Apr0088. The 50L Demonstration (C235 MCB) BLI titer was determined using KBI Reference Material lot # S-20210314-0001-SD2-E-M (derived from Process Feasibility Run #2, Section 3.3.1). The 50L GMP run (C235 MCB) BLI titer was determined using KBI Reference Material lot # P65. Although absolute titers varied across runs (mainly due to different reference standards), studies demonstrated robust production and high recovery efficiency under the same feed and control strategy.

In summary, the 50 L material Supply (RCB), 50 L demonstration (MCB), and 50 L GMP runs confirm that the fed-batch process developed at the ambr250 scale is scalable and reproducible across both cell bank sources and vessel scales (XDR-50 and XDR-200). Similar cell growth, metabolite behavior and BLI titers, along with high harvest yields in both runs, demonstrate that the process parameters were suitable for clinical manufacturing.

### 3.3 Downstream Purification Process development

#### 3.3.1 Initial Process Evaluation during Feasibility Runs at Bench Scale

The initial process evaluations included the feasibility run and robustness. As shown in Table 7, the step yields for center point cycles of each unit operation were in line with expectations. For the robustness evaluation, when operating 2G12 capture and MabSelect SuRe chromatography unit operations, within theoretical worst-case yield conditions, the yields observed were acceptable. For 2G12, the range observed for center point cycles was 58.9±1.3% and during the worst-case yield execution, the step yield was still 58%. For MSS, the step yield observed for the center point cycle was 98% compared to 95% during the worst-case yield execution.

**Table 7:** Feasibility Run and Robustness– Operational Parameters and Process Performance.

| Unit Operation | Cycle | Column Loading (g/L<sub>resin</sub>) | Yield | Study |
| --- | --- | --- | --- | --- |
| Capture (2G12) | 1-3 | 1.30 | 58.9±1.3 | Centerpoint |
|  | 4 | 1.30 | 58.7% | WCPQ |
|  | 5 | 0.75 | 57.7% | WCY |
| UF/DF1 | N/A | 29 g/m<sup>2</sup> | 85.4% | Centerpoint |
| Triton X-100 VI & Amberlite Adsorption | N/A | N/A | 91.9% | Centerpoint |
| MabSelect SuRe (flowthrough) | 1 | 5.00 | 94.6% | WCY |
|  | 2 | 20.04 | 97.7% | Centerpoint |
| Capto adhere (flowthrough) | 1 | 2.13 | 23.9% | WCY |
|  | 2 | 5.00 | 60.8% | WCPQ |
| Planova 20N VF | N/A | 39.7 L/m<sup>2</sup> | 95.4% | Fresh (non-frozen load) |
|  |  | 67.2 L/m<sup>2</sup> | 96.6% | Fresh (non-frozen load) |
|  |  | 67.1 L/m<sup>2</sup> | 96.4% | Freeze / thawed load |
| UF2 | N/A | N/A | 84.1% | Centerpoint |
| Superdex 200pg | 1 | 1.0% CV | 80.5% | WCY |
|  | 2-4,6-7 | 2.0% CV | 89.4±1.0 | Centerpoint |
|  | 5 | 1.0% CV | 78.4% | WCY |

Alternatively, for the Capto adhere robustness evaluations, it was observed that the yield decreased from ∼60% to approximately 24% when the step was operated under theoretical worst-case yield conditions. As a result, for cGMP manufacturing it was recommended to monitor the loading factor and pH during load adjustment for Capto adhere such that conditions near worst-case yield are avoided where at all possible.

The product quality results for the initial process evaluation are presented in Table 8 and Table 9.

**Table 8:** Feasibility Run and Robustness– SE-HPLC and RP-HPLC Results.

| Unit Operation | Intermediate | Cycle | SE-HPLC |  |  | RP-HPLC |  |  | Study |
| --- | --- | --- | --- | --- | --- | --- | --- | --- | --- |
|  |  |  | HMW (%) | Main (%) | LMW (%) | Pre-Main (%) | Main (%) | Post-Main (%) |  |
| Capture (2G12) | Diluted Eluate | 1-3 | 9.7±0.9 | 74.8±4.4 | 15.5±3.7 | 0.1± 0.1 | 74.8±2.8 | 25.2±2.8 | Centerpoint |
|  |  | 4 | 10.0 | 80.8 | 9.2 | 0.0 | 75.7 | 24.3 | WCPQ |
|  |  | 5 | 9.0 | 77.4 | 13.6 | 0.0 | 74.6 | 25.4 | WCY |
| UF/DF1 | Retentate | N/A | 13.1 | 85.5 | 1.4 | 0.0 | 75.4 | 24.6 | Centerpoint |
| Triton VI / Amberlite | Flowthrough | N/A | 11.3 | 87.3 | 1.3 | 0.0 | 73.1 | 27.0 | Centerpoint |
| MabSelect SuRe | Flowthrough | 1 | 10.7 | 88.0 | 1.3 |  |  |  | WCY |
|  |  | 2 | 10.9 | 88.0 | 1.1 |  |  |  | WCPQ |
| Capto adhere | Load | 1 | 10.6 | 88.2 | 1.2 |  |  |  | WCY |
|  | Flowthrough |  | 0.3 | 98.4 | 1.4 | 0.0 | 86.5 | 13.5 |  |
|  | Load | 2 | 10.7 | 88.3 | 1.1 |  |  |  | WCPQ |
|  | Flowthrough |  | 0.4 | 98.5 | 1.2 | 0.0 | 83.2 | 16.8 |  |
| Planova 20N VF | Filtrate | N/A | 0.5 | 98.3 | 1.2 |  |  |  |  |
| UF2 | Retentate | N/A | 0.4 | 98.9 | 0.7 |  |  |  | Centerpoint |
| Superdex 200pg | Flowthrough | 1 | 0.3 | 98.8 | 0.9 |  |  |  | WCY |
|  |  | 2-4,6-7 | 0.3± 0.0 | 98.9±0.1 | 0.8±0.1 |  |  |  | Centerpoint |
|  |  | 5 | 0.3 | 98.8 | 0.9 |  |  |  | WCY |
|  |  | Cycle 1-4 Pool | 0.1 | 98.9 | 1.0 |  |  |  |  |
|  |  | Cycle 5-7 Pool | 0.1 | 99.4 | 0.5 | 0.0 | 85.4 | 14.5 |  |

**Table 9:** Feasibility Run and Robustness– Residual Impurity Clearance and BLI Relative Potency.

| Unit Operation | Intermediate | Cycle | resHCP (ppm) | resDNA (ppb) | resProA (ppm) | resTriton X-100 | res2G12 (ppm) | BLI Relative Potency | Study |
| --- | --- | --- | --- | --- | --- | --- | --- | --- | --- |
| Capture (2G12) | Diluted Eluate | 1-3 | 15521±578 | 3401±70 |  |  |  |  | Centerpoint |
|  |  | 4 | 14575 | 3756 |  |  |  |  | WCPQ |
|  |  | 5 | 11668 | 1499 |  |  |  |  | WCY |
| UF/DF1 | Retentate | N/A | 27845 |  |  |  |  |  | Centerpoint |
| Triton VI / Amberlite | Flowthrough | N/A | 20116 |  |  | Below LOQ (< 0.002%) |  |  | Centerpoint |
| MabSelect SuRe | Flowthrough | 1 | 21966 |  | 155 |  |  |  | WCY |
|  |  | 2 | 16065 |  | Below LOQ (< 48 ppm) |  |  |  | WCPQ |
| Capto adhere | Load | 1 | 18807 | 2499 | - |  |  |  | WCY |
|  | Flowthrough |  | Below LOQ (< 63 ppm) | Below LOQ (< 27 ppb) | Below LOQ (< 2 ppm) |  |  |  |  |
|  | Load | 2 | 11287 | 2810 | - |  |  |  | WCPQ |
|  | Flowthrough |  | 296 | 105 | Below LOQ (<1 ppm) |  |  |  |  |
| Planova 20N VF | Filtrate | N/A | 228 |  |  |  |  |  |  |
| UF2 | Retentate | N/A | 34 |  |  |  |  |  | Centerpoint |
| Superdex 200pg | Flowthrough | Cycle 1-4 Pool | Below LOQ (< 11.4 ppm) |  |  |  |  |  | WCY |
|  |  | Cycle 5-7 Pool | Below LOQ (< 12.2 ppm) | Below LOQ (< 6 ppb) | Below LOQ (< 1 ppm) | Below LOQ (< 0.002%) | Below LOQ (< 30 ppm) | 100% | Centerpoint/ WCY pool |

By SE-HPLC, the 2G12 capture eluate for the centerpoint runs contained approximately 9.7±0.9% HMW and 15.5±3.7% LMW, however the process was able to reduce both HMW and LMW to < 1.0% in the final material. The clearance of these species increased the % Main product from approximately 74.8±4.4% in the 2G12 capture eluate to > 99% in the final material. Additionally, by RP-HPLC the % Main species was increased throughout the process, with the undesired % Post-Main species reduced to acceptable levels.

For the robustness evaluation, the 2G12 elute from the worst-case product quality run contained approximately 81% Main species SE-HPLC and 76% Main species by RP-HPLC, with profiles comparable to the centerpoint runs. For Capto adhere, the flowthrough from the worst-case product quality run contained approximately 99% Main species SE-HPLC and 83% Main species by RP-HPLC, with profiles comparable to the centerpoint run. The observation demonstrated the robustness of these two-unit operations with respect to product-related impurities.

Table 9 Contains the residual impurity clearance results, including resHCP, resDNA, resProA, resTriton X-100, and res2G12, as well as BLI relative potency. The downstream process demonstrated the ability to remove each residual impurity so that all final targets were met. The final purified preparative SEC eluate was used as the reference material for the BLI relative potency assay. For the robustness evaluations, the 2G12 elute from the worst-case product quality run contained approximately 15,000 ppm resHCP and 3,800 ppb resDNA, which was comparable to the centerpoint runs. For Capto adhere, the flowthrough from the worst-case product quality run contained approximately 300 ppm resHCP, 100 ppb resDNA, <LOQ, and 1 ppm resProA. Although the resHCP and resDNA were 3- to 4-fold higher in this eluate compared with the centerpoint condition, the impurity levels were considered acceptable for this unit operation since further downstream unit operations have demonstrated the ability to remove remaining amounts of these impurities. Capto adhere mixed-mode chromatography achieves substantial reduction of residual impurities within the N332-GT-5 gp140 downstream process even under worst-case product quality conditions. This aligns with the application of mixed-mode resins to reduce residual impurities, as reported by Wolfe et al., 2021, for complex glycoproteins.

#### 3.3.2 Evaluation of Removal of Preparative SEC Chromatography

For the clinical cGMP supply run, there was a procurement challenge for Superdex 200pg resin. The biopharmaceutical industry, in general, faced major supply chain disruption at the time of process development of N332-GT5 gp140 (Socal et al., 2021). An evaluation was completed to assess the performance of a final UF/DF without a preparative SEC step, where conditions from UF2 and UF/DF 3 were combined into a single operation. This execution was compared to an execution using the original process containing SEC. The two executions were performed in parallel. Table 10 shows that the final UF/DF step was able to remove impurities from the product pool to an acceptable level and the overall results were comparable to the control execution with preparative SEC. Since the product quality (SE-HPLC and RP-HPLC) and residual HCP levels are comparable, Superdex 200pg chromatography unit operation was removed from the process for cGMP run.

**Table 10:** Preparative SEC Removal Study – Product Quality.

| Experiment | Unit Operation | Intermediate | resHCP (ppm) | SEC-HPLC (LMW) | RP-HPLC (pre-Main) |
| --- | --- | --- | --- | --- | --- |
| Control | Superdex 200pg | Load | < LOQ<br>(< 157 ng/mg) | 1.8% | 1.5% |
|  |  | Flowthrough | < LOQ<br>(< 34 ng/mg) | 0.8% | 0.5% |
| Preparative SEC Removal | Final UF/DF | Load | < LOQ<br>(< 157 ng/mg) | 1.8% | 1.5% |
|  |  | Final Retentate | < LOQ<br>(< 29 ng/mg) | 1.0% | 0.8% |

#### 3.3.3 Intermediate Hold Time Stability Study

Downstream process intermediates were held for 3 days at ambient temperature and 7 days at 2–8 °C, then evaluated by SE-HPLC and relative potency. Figure 15 shows that a 3-4% increase in HMW was observed in the UF/DF1 retentate from Day 0 to Day 1 at both ambient and 2–8 °C, prompting a ≤1-day hold time recommendation for this intermediate across both temperature ranges. The % HMW increased by 0.7% in VIN retentate from Day 2 to Day 3 at ambient temperature, so out of caution, a ≤ 2-day hold was recommended at ambient temperature for VIN retentate. A decreasing trend in % LMW was observed for the 2G12 diluted eluate over the hold times at both temperatures, but a decrease in impurities is not considered an at-risk change, so these data did not impact the recommended hold time. All other process intermediates showed ≤ 0.7% change in HMW and LMW species over the tested time frames by SE-HPLC. Lastly, a decreasing trend in relative potency of the BDS was observed over the evaluated hold times, but it was still within the target potency range for the BDS, and the SE-HPLC data were not impacted; therefore, the recommendation was not to alter the hold time based on these results. Combining these data, all intermediates except the UF/DF 1 retentate and the VIN eluate were assigned a recommended hold time of up to 3 days at ambient temperature and 7 days at 2–8 °C, since there was no observed impact on product quality at the maximum duration evaluated.

**Figure 15:**
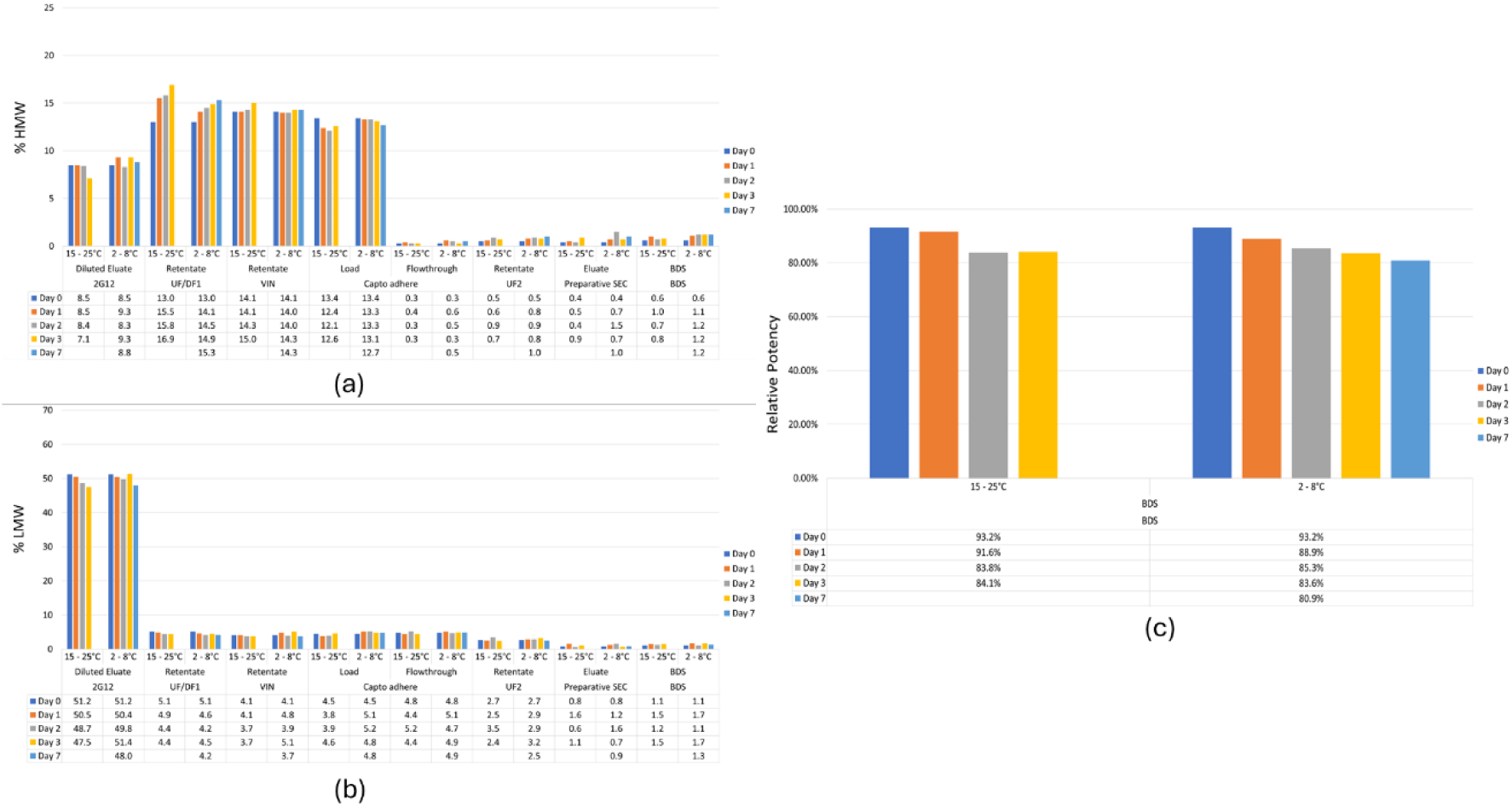
Intermediate hold time stability study: An intermediate hold time study was conducted using pilot-scale material. Depicted are (a) % HMW by SE-HPLC, (b) % LMW by SE-HPLC and (c) Relative potency at BDS

### 3.4 Pilot-scale and cGMP Downstream Processing of Drug Substance

The step yields from the pilot-scale demonstration run are shown in Figure 16. Overall downstream process yield was 48.8%. The observed >100% yield for 2G12 step can likely be attributed to the titer assay variability associated with clarified harvest generated from the upstream demonstration run. Differences in the step yield for Capto adhere are not unexpected based on the results from development during the feasibility run and robustness evaluations. With respect to product quality, the downstream process reduced %HMW levels to < 1%, with % Main Peak > 98% (SE-HPLC), which is comparable to small-scale data. All residual impurities were reduced to <LOQ, with Capto adhere providing the majority of the clearance, also consistent with small-scale data.

**Figure 16:**
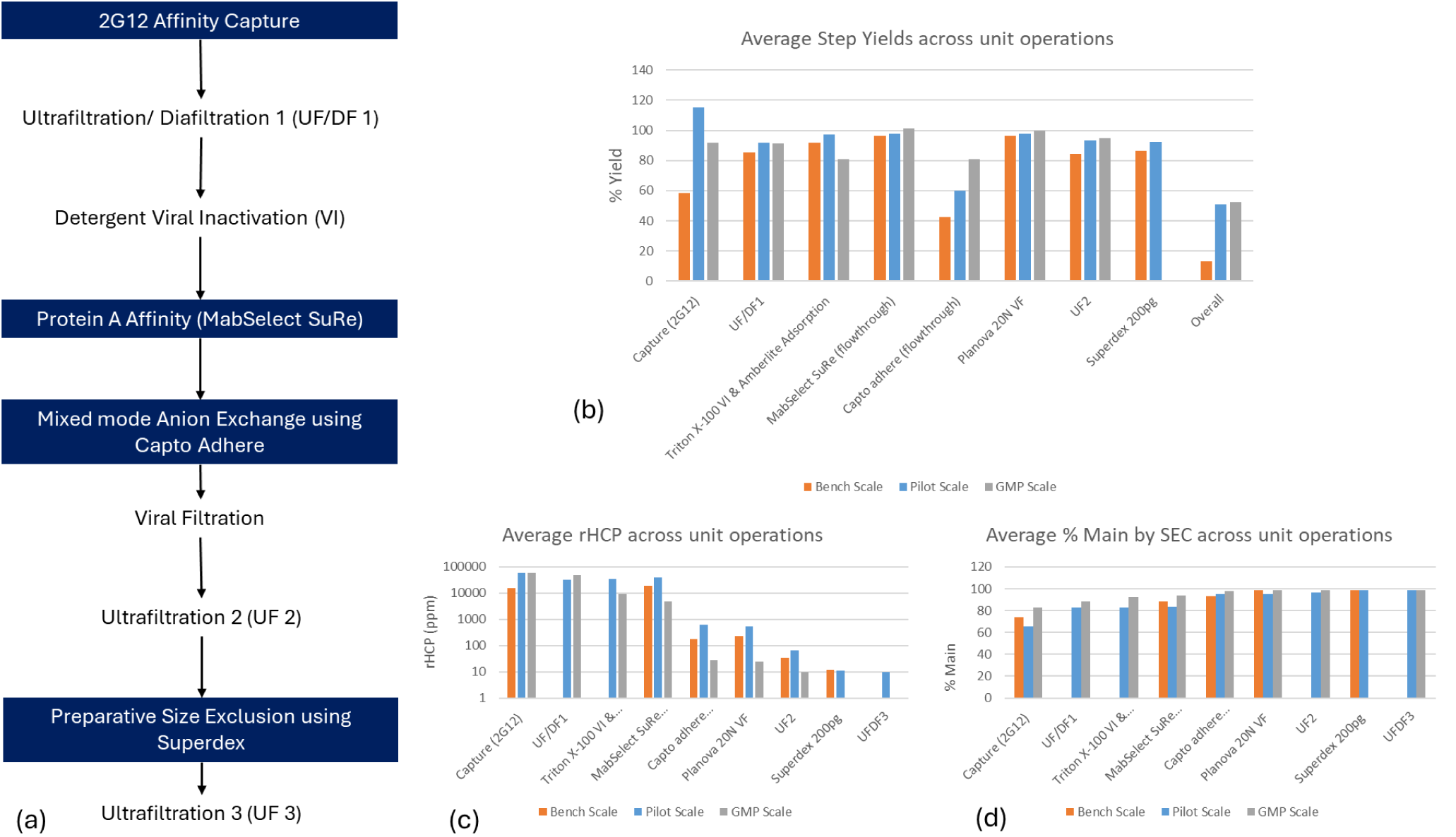
Downstream Process and Performance. Demonstration run pilot-scale utilized 10 – 20 cm ID columns. cGMP scale utilized 14 – 45 cm ID columns. Depicted are (a) Process overview, (b) Average step yields for each unit operation, (c) ResHCP levels of each process intermediate, (d) % Main peak by SE-HPLC.

The step yields, SE-HPLC and residual impurity levels observed during the cGMP run are also shown in Figure 16. As discussed in section 3.3.2, the preparative SEC step was not performed at cGMP scale due to unavailability of the resin and studies supporting removal of the step from the process. The step yields were again comparable to small- and pilot-scale, with the exception of the 2G12 capture and Capto adhere which were expected to have variability based on the prior development data. Table 11 shows the product quality comparison for BDS from the pilot-scale and cGMP scale. The data are comparable for all assays, indicating similar performance and scalability of the downstream process.

**Table 11:** Comparison of Bulk Drug Substance product quality for the non-GMP batch (pilot-scale) and cGMP Batch production runs of N332-GT5 gp140 trimer.

| Attribute | Pilot-scale Demonstration Batch | GMP Production |
| --- | --- | --- |
| Appearance | Clear colorless liquid, free of visible particulates | Colorless solution |
| pH | 7.5 | 7.5 |
| Concentration | 1.0 mg/mL | 1.0 mg/mL |
| Purity (RP-HPLC) | Comparable to Reference | Comparable to Reference |
| Percent Trimer (SE-HPLC) | %Main Peak: 98.6%<br>%HMW: 0.2%<br>%LMW: 1.2% | %Main Peak: 99.4%<br>%HMW: N/A<br>%LMW: 0.6% |
| Percent trimer (BG18_GL0 BLI) | NT | 116% Compared to Reference Standard |
| Residual CHO HCP | < LOQ<br>(< 10 ng/mg) | < LOQ (42 ng/mg) |
| Residual CHO DNA | < LOQ<br>(< 4.5 ppb) | < LOQ of 5 pg/mg |
| Residual Protein A | < LOQ<br>(< 0.3 ng/mg) | < LOQ of 1 ng/mg |
| Residual 2G12 | < LOQ<br>(< 25 ng/mg) | < LOQ of 12.5 ng/mg |
| Bioburden | 0 CFU/10mL | 0 CFU/10 mL |
| Endotoxin | 0.1503 EU/mL | < 0.01 EU/mL |

### 3.5 Virus Clearance Study

A common industry standard for retroviral log reduction is to achieve a reduction of at least four logs beyond the estimated viral load per therapeutic dose, which is determined using retrovirus-like particle (RVLP) counts in the bioreactor supernatant (Shukla & Aranha, 2015). As shown in Table 12, for the downstream process for N332-GT5 gp140 the total log reduction achieved for XMuLV was ≥ 18.14 logs, indicating that the expected retroviral load per dose would be ≤ 4.0 x 10⁻⁹ RVLP. This corresponds to less than one viral particle per 250 million doses and greatly surpasses the industry benchmark of ≤ 10⁻⁴ RVLP per dose. For parvovirus removal, the standards are to have a unit operation with at least 4 LRV and a minimum of two orthogonal steps providing clearance. For MMV, the total process log reduction was ≥ 11.70 logs and includes three distinct unit operations. These results demonstrate that the N332-GT5 gp140 purification unit operations can effectively achieve the target viral clearance, significantly reducing the risk of viral contamination in the final product.

**Table 12:** Virus Removal/Inactivation of XMuLV and MMV by purification process for N332-GT5 gp140.

| Purification Process step | Log<sub>10</sub> Virus Reduction |  |
| --- | --- | --- |
|  | XMuLV | MMV |
| 2G12 Affinity Chromatography | 2.95 ± 0.57 | 2.19 ± 0.49 |
| Triton X-100 Detergent Inactivation | 3.91 ± 0.17 | NA |
| Capto Adhere Chromatography | $\geq 6.11 \pm 0.34$ | $\geq 6.19 \pm 0.25$ |
| Viral Filtration (Planova 20N) | $\geq 5.17 \pm 0.31$ | $3.32 \pm 0.27$ |
| **Overall** | **$\geq 18.14 \pm 0.75$** | **$\geq 11.70 \pm 0.61$** |

### 3.6 Product Characterization

#### 3.6.1 Analysis of Site-specific N-glycosylation by Mass Spectrometry

Two complementary mass spectrometry methods were used to analyze the glycosylation at the 27 N-glycosylation sites (PNGS) in the Env glycoprotein (Baboo et al., 2021; Cao et al., 2017; Behrens et al., 2017; Go et al., 2017; Watanabe et al., 2020). Combined, the two methods show high glycan occupancy at most sites and establish the extent to which the immature oligomannose-type glycan is trimmed and converted to complex type glycans with terminal sugars including sialic acid.

The first method, DeGlyPHER (Baboo et al., 2021), is a proteomics platform that generates glycopeptides using a non-specific protease, followed by sequential treatment with two glycosidases that cleave oligomannose-type glycans (Endo H) and complex-type glycans (PNGase F), leaving mass signatures on the asparagine of +203 and +3, respectively. Then, using standard MS/MS proteomics each glyco-peptide can be assessed for carrying no-glycan (+0), immature oligomannose (+203), or mature complex glycan (+3). The method is highly sensitive, enabling quantitative analysis of the glycosylation status at each PNGS. Results using this method (Figure 17) show that most PNGS are fully occupied with glycan (>95%), four PNGS (N185e, N185h, N611, N618) are partially unoccupied (10-20%), and only one PNGS (N625) is substantially unoccupied (50%). Also evident is that most of the PNGS are predominantly populated with oligomannose-type glycans, reflecting minimal glycan processing, while glycans at five PNGS are substantially processed to the mature complex type glycans (N185e, 185h, N462, N611, N618).

**Figure 17:**
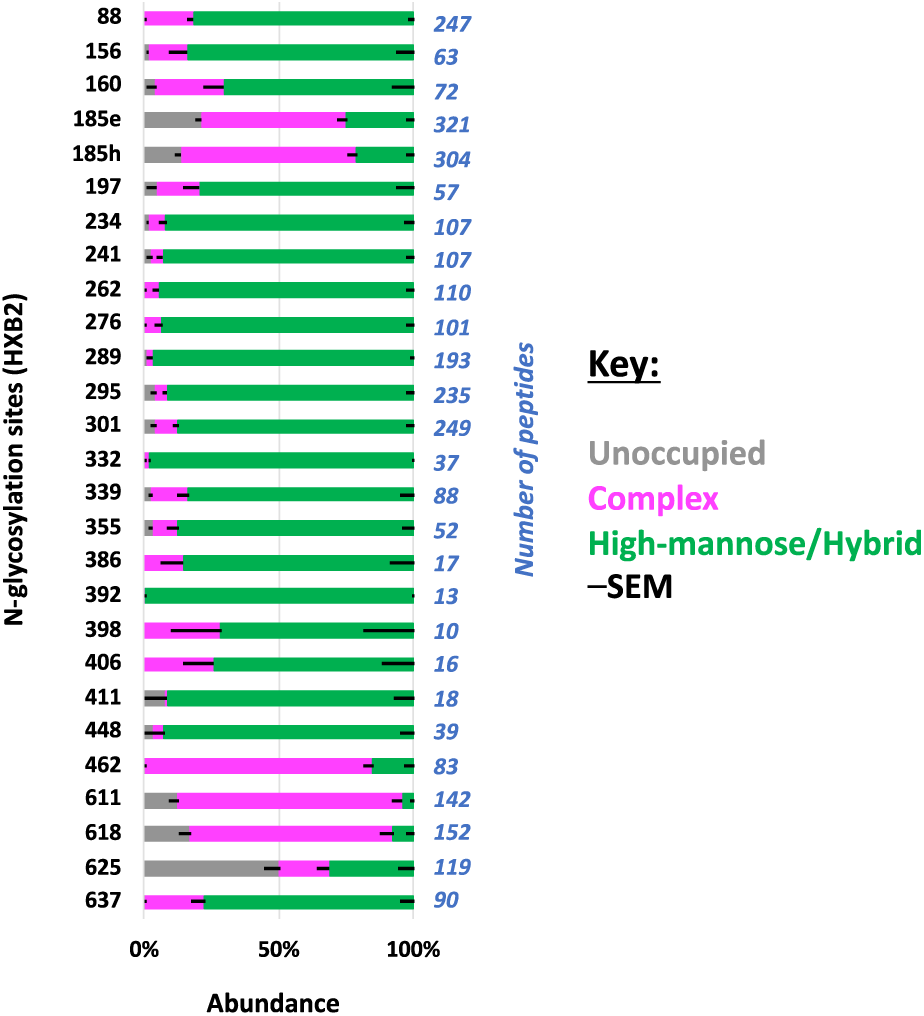
Site-specific N-glycosylation analysis using DeGlyPHER. Oligomannose-type glycans are shown in green, complex and hybrid-type glycans in magenta, and unoccupied sites in grey. Data are shown as the mean of abundance measurement of peptides mapping to individual PNGS with the standard error of mean.

The complementary method employs traditional liquid chromatography-mass spectrometry (LC-MS) glycoproteomics using several specific proteases with glycopeptides of known sequence retaining intact N-glycans (Behrens et al., 2017). The MS data from glycopeptides are searched for known peptides without modification (no occupancy) and with modification by glycans of known molecular weight from a glycoform library of candidate oligomannose- and complex-type glycans. As shown in Figure 18, results for each PNGS display the glycoforms identified. Because this method relies on the ionization of particular peptides and glycopeptides containing a single glycosylation site, not all PNGS are represented. However, for PNGS with glycopeptides detected there is good agreement with the results in Figure 17 as to which sites retain oligomannose-type glycans with minimal processing, and which sites are substantially processed to complex glycoforms. Most sites that are predominantly oligomannose-type (e.g. N262, N332) have some trimming of the Man_9_GlcNAc_2_ glycoform to smaller glycoforms, which is the information provided uniquely by this method. Notably, at several PNGS only the non-occupied peptides were prominent suggesting the absence of glycan at that position (N185e, N611, N625). However, these sites are shown to be 50-90% glycosylated by the other method (Figure 17), revealing that aassessment of occupancy exhibits higher variation between analytical approaches.

**Figure 18:**
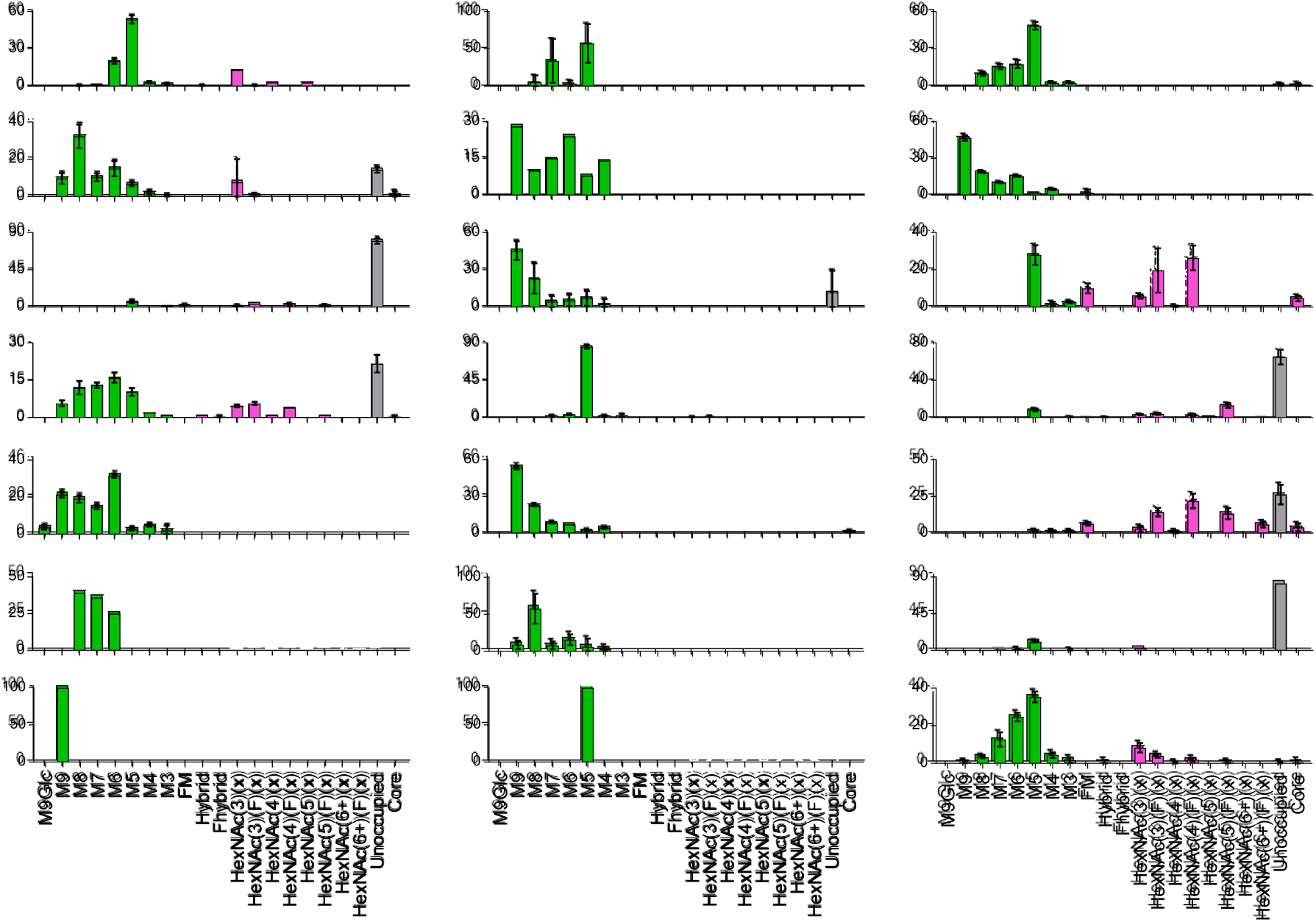
Site-specific N-glycosylation analysis using LC-MS glycoproteomics. - Site-specific glycan content was determined by LC-MS on intact glycopeptides. The average (+/- SEM) proportion of each type of glycan is depicted on the y-axes (%), the glycan type on the horizontal category axis. Oligomannose-type glycans are displayed green, complex-type glycans, magenta and the proportion of unoccupied in grey.

#### 3.6.2 Negative-stain Electron Microscopy

Single-particle negative-stain electron microscopy (nsEM) was used to assess the structural integrity of N332-GT5 gp140 from the demonstration run. Micrographs revealed monodisperse protein particles, with no evidence of aggregation or visible impurities (Figure 19). 2D class averages of the imaged particles demonstrate that all particles have a similar phenotype of three, compact lobes, indicative of native-like Env trimers (Figure 19). No other oligomeric states (dimers, monomers) were observed in the nsEM analysis, confirming the presence of nearly 100% native-like trimers, similar to published cGMP production runs of other engineered Env trimers (Dey et al., 2018; Bale et al., 2025).

**Figure 19:**
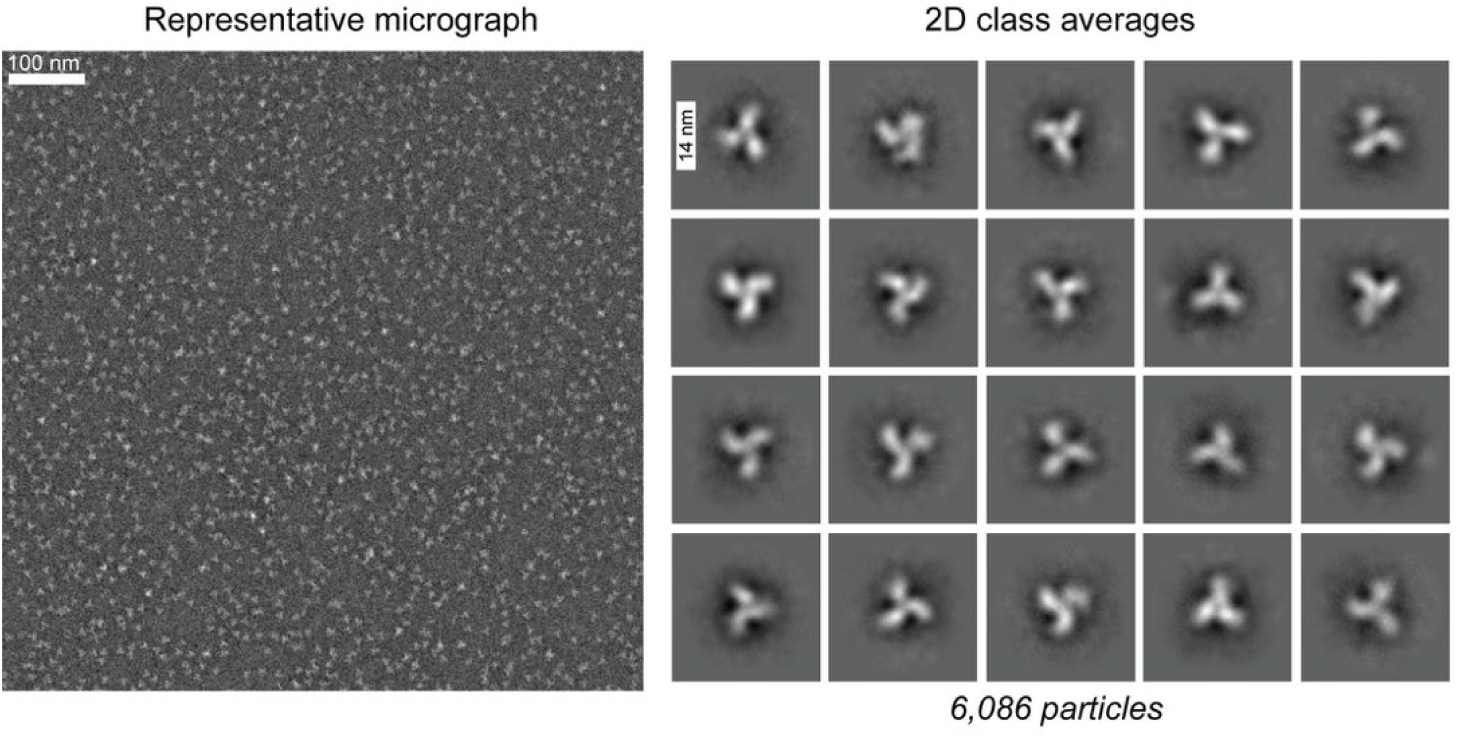
Negative-stain electron microscopy images of N332-GT5 gp140 demonstration run. Representative micrograph (*left*) and 2D class averages (*right)* with respective scale bars.

## 4 Conclusions

We established a stable CHO cell-based platform for N332-GT5 gp140 manufacturing using Leap-In transposition, followed by rational clone selection and rigorous genetic stability qualification. The upstream process scaled seamlessly from Ambr^®^ 250 to 200 L SUBs; downstream processing produced >99% trimer with excellent antigenic integrity and defined glycosylation. Intermediate hold-time stability data confirmed that the process produces product that is stable under manufacturing operations. Viral clearance far exceeded benchmarks for both enveloped and non-enveloped models, and hold-time studies defined operationally practical limits without compromising quality. Together, these data demonstrate the feasibility of reliable, clinical-grade production of native-like HIV-1 Env trimers and provide a blueprint for advancing glycoprotein vaccine candidates through early clinical phases. Similar approaches for other candidate molecules should help shorten development timelines leading up to clinical trials and are well-suited for HIV-1 Env trimer process development.

## Contributions

S.P., A.E.P., K.S., and D.C. developed the SoW and oversaw the development, GMP manufacturing studies, and Data analysis.

S.U, T.B, A.C., J.W., and L.W – executed the upstream, downstream development studies, and analytical development

F.C and V.S – Executed the Cell line development studies

W-H.L. and G.O. performed the EM analysis and determined the cryo-EM structure. A.B.W. oversaw the structural analysis

S.B, J.D.A - performed glycan analysis. J.R.Y, J.C.P and M.C. supervised the glycan analysis J.S and W.R.S - designed the trimer construct

S.P, S.U, F.B, S.B, J.CP, G.O – Authored the paper

All authors reviewed the results. All authors read and approved the final version of the manuscript.

Corresponding authors: Correspondence to Sammaiah Pallerla or Shaunak Uplekar

## References

Baboo S, Diedrich JK, Martínez-Bartolomé S, Wang X, Schiffner T, Groschel B, et al. DeGlyPHER: An ultrasensitive method for the analysis of viral spike N-glycoforms. Analytical Chemistry. 2021;93(40):13651–13657. 10.1021/acs.analchem.1c03059

Baboo S, Diedrich JK, Martínez-Bartolomé S, Wang X, Schiffner T, Groschel B, et al. DeGlyPHER: Highly sensitive site-specific analysis of N-linked glycans on proteins. Methods in Enzymology. 2023;682:137–185.

Bale S, et al. Accelerated cGMP production of near-native HIV-1 Env trimers following electroporation transfection and immunogenicity analysis. npj Vaccines. 2025.

Behrens AJ, Harvey DJ, Milne E, Cupo A, Kumar A, Zitzmann N, et al. Molecular architecture of the cleavage-dependent mannose patch on a soluble HIV-1 envelope glycoprotein trimer. Journal of Virology. 2017;91(2):e01894–16. 10.1128/JVI.01894-16

Burton DR, Hangartner L.Broadly neutralizing antibodies to HIV and their role in vaccine design.Annual Review of Immunology. 2016;34:635–659.

Cao L, Diedrich JK, Kulp DW, Pauthner M, He L, Park SR, et al. Global site-specific N-glycosylation analysis of HIV envelope glycoprotein. Nature Communications. 2017;8:14954.

Dey AK, Cupo A, Ozorowski G, Sharma VK, Behrens AJ, Go EP, et al. cGMP production and analysis of BG505 SOSIP.664, an extensively glycosylated trimeric HIV-1 envelope glycoprotein vaccine candidate. Biotechnology and Bioengineering. 2018;115:885–899. 10.1002/bit.26498

Go EP, Ding H, Zhang S, Ringe RP, Nicely N, Hua D, et al.Glycosylation benchmark profile for HIV-1 envelope glycoprotein production based on eleven Env trimers. Journal of Virology. 2017;91(9):e02428–16.

Hahn WO, Parks KR, Shen M, Ozorowski G, Janes H, Ballweber-Fleming L, et al. Use of 3M-052-AF with alum adjuvant in HIV trimer vaccine induces human autologous neutralizing antibodies. Journal of Experimental Medicine. 2024;221(10):e20240604. 10.1084/jem.20240604

HIV Vaccine Trials Network. HVTN 144: A Phase 1 clinical trial to evaluate the safety and immunogenicity of N332-GT5 gp140 adjuvanted with SMNP. ClinicalTrials.gov Identifier: NCT05217641.

Klasse PJ, Moore JP.Antibodies to the envelope glycoproteins of HIV-1: structure, function, and mechanisms of action. Progress in Lipid Research. 2012;51:253–269.

Kwong PD, Mascola JR. HIV-1 vaccines based on antibody identification, B-cell ontogeny, and epitope structure. Immunity. 2018;48:855–871.

Pallerla S, et al.,. Scale-up and cGMP manufacturing of next-generation vaccine adjuvant saponin/MPLA nanoparticles (SMNP). Journal of Pharmaceutical Sciences. 2025. 10.1016/j.xphs.2025.103913

Park SK, Venable JD, Xu T, Yates JR.A quantitative analysis software tool for mass spectrometry-based proteomics. Nature Methods. 2008;5:319–322.

Pauthner M, Havenar-Daughton C, Sok D, et al.Elicitation of robust Tier-2 neutralizing antibody responses in nonhuman primates by HIV envelope trimer immunization using optimized approaches. Immunity. 2017;46:1073–1088.

Peng J, Elias JE, Thoreen CC, Licklider LJ, Gygi SP. Evaluation of multidimensional chromatography coupled with tandem mass spectrometry for large-scale protein analysis: the yeast proteome.Journal of Proteome Research. 2003;2:43–50.

Rameez S, Mostafa S, Miller C, Shukla A. High-throughput miniaturized bioreactors for cell culture process development: reproducibility, scalability, and control. Biotechnology Progress. 2014;30:718–727.

Ramezani-Rad P, Cottrell CA, Marina-Zárate E, et al.Vaccination with an mRNA-encoded membrane-bound HIV envelope trimer induces neutralizing antibodies in animal models. Science Translational Medicine. 2025;17(809):adw0721. 10.1126/scitranslmed.adw0721

Sanders RW, Derking R, Cupo A, Julien JP, Yasmeen A, de Val N, et al. A next-generation cleaved, soluble HIV-1 Env trimer, BG505 SOSIP.664 gp140, expresses multiple epitopes for broadly neutralizing but not non-neutralizing antibodies. PLoS Pathogens. 2013;9:e1003618.

Shukla A, Aranha H.Viral clearance for biopharmaceutical downstream processes. Pharmaceutical Bioprocessing. 2015;3:127–138.

Silva M, et al. A stabilized HIV-1 envelope trimer that mimics the native viral spike induces neutralizing antibodies.Science Immunology. 2021;6:eabf1152.

Socal M, Sharfstein J, Greene J. The pandemic and the supply chain: gaps in pharmaceutical production and distribution. American Journal of Public Health. 2021;111:635–639.

Steichen JM, et al.Vaccine priming of rare HIV broadly neutralizing antibody precursors in nonhuman primates. Science. 2024;384:adj8321.

Steichen JM, Lin YC, Havenar-Daughton C, et al.A generalized HIV vaccine design strategy for priming of broadly neutralizing antibody responses.Science. 2019;366:eaax4380.

Suloway C, Pulokas J, Fellmann D, Cheng A, Guerra F, et al.Automated molecular microscopy: the new Leginon system. Journal of Structural Biology. 2005;151:41–60.

Tabb DL, McDonald WH, Yates JR. DTASelect and Contrast: tools for assembling and comparing protein identifications from shotgun proteomics. Journal of Proteome Research. 2002;1:21–26.

UNAIDS. Global HIV & AIDS statistics — 2025 fact sheet.https://www.unaids.org

Watanabe Y, Berndsen ZT, Raghwani J, Seabright GE, Allen JD, Pybus OG, et al. Vulnerabilities in coronavirus glycan shields despite extensive glycosylation. Nature Communications. 2020;11:2688.

Wolfe L, Smedley J, Bubna N, Hussain A, Harper R, et al. Development of a platform-based approach for the clinical production of HIV gp120 envelope glycoprotein vaccine candidates. Vaccine. 2021;39:3852–3861.

Xu T, Park SK, Venable JD, Wohlschlegel JA, Diedrich JK, Cociorva D, et al. ProLuCID: an improved SEQUEST-like algorithm with enhanced sensitivity and specificity. Journal of Proteomics. 2015;129:16–24.

Steichen, et al., HIV Vaccine Design to Target Germline Precursors of Glycan-Dependent Broadly Neutralizing Antibodies, Immunity, 2016. Volume 45, Issue 3 p483–496.

Parks, et al., Vaccination with mRNA-encoded membrane-anchored HIV envelope trimers elicited tier 2 neutralizing antibodies in a phase 1 clinical trial. Science Translational Medicine, 30 Jul 2025, Vol 17, Issue 809. DOI: 0.1126/scitranslmed.ady6831

Sander, et al. HIV-1 neutralizing antibodies induced by native-like envelope trimers. Science, Vol 349, Issue 6244. DOI: 10.1126/science.aac4223

